# Fluctuations in functional connectivity associated with interictal epileptiform discharges (IEDs) in intracranial EEG

**DOI:** 10.1101/2021.05.14.444176

**Authors:** Jennifer Stiso, Lorenzo Caciagli, Peter Hadar, Kathryn A. Davis, Timothy H. Lucas, Dani S. Bassett

## Abstract

All epilepsies are defined by a propensity for recurrent seizures, characterized by hypersynchronous electrographic activity. Understanding this overarching property would be advanced by a thorough quantification of how the global synchrony of the epileptic brain responds to small perturbations that do not trigger seizures. Here, we leverage analysis of transient focal bursts of epileptiform activity, termed interictal epileptiform discharges (IEDs), to characterize this response. Specifically, we use a group of 145 participants implanted with intracranial EEG (iEEG) electrodes to quantify changes in five functional connectivity measures associated with three properties of IEDs: their presence, spread, and number. We perform this analysis in five frequency bands in order to contextualize our findings in relation to ongoing neural processes at different spatial and temporal scales. We find that, across frequency bands, both the presence and spread of IEDs tend to lead to independent increases of functional connectivity, but only in functional connectivity measures influenced by the amplitude, rather than the phase, of a signal. We find that these increases are not explained by simple subgroups of connections, such as the weakest connections in the brain, or only connections within the seizure onset zone. Evaluating patterns of similarity across different bands and measure combinations, we find that the presence of IEDs impacts high frequencies (gamma and high gamma) and low frequencies (theta, alpha, and beta) differently, although responses within each group are similar. Using grouped LASSO regression, we identify which individual-level features explain differences in functional connectivity changes associated with IEDs. While no single feature robustly explains observed differences, the most consistently included predictors across bands and measures are the rate of IEDs and the anatomical locus of IEDs. Overall, this work provides compelling evidence for increases in global synchrony associated with IEDs, and delivers a thorough exploration of different functional connectivity measures, frequency bands, and IED properties. These observations show a disruption of several types of ongoing neural dynamics associated with IEDs. Additionally, we provide a starting point for future models of how small perturbations affect neural systems and how those systems support the hypersynchrony seen in epilepsy.

## INTRODUCTION

Epilepsy is a heterogeneous neurological disease that is characterized by a predisposition towards recurrent seizures[1, 2]. While the different subtypes of epilepsy manifest in different types of seizures, levels of severity, and cognitive or somatic comorbidities, the macroscale behavior of the brain exhibits a marked tendency towards hypersynchronous activity evident in seizures[1, 3]. In order to better understand this property of hypersynchrony, it is useful to investigate how small perturbations affect endogenous patterns of neural synchrony. An interictal epileptiform discharge (IED) is a sub-seizure epileptic waveform that can either trigger or prevent the broad hypersynchrony characteristic of seizure dynamics[4, 5]. Similar to the diverse phenotypes of epilepsy, IEDs also have diverse waveforms, and are generated by several distinct, overlapping functional networks[6, 7]. This heterogeneity results in different sequences of IEDs having different spatiotemporal properties within an individual. However, across these different functional groupings, IEDs consistently precede seizure activity and occur in spatially proximal regions [8]. Therefore, IEDs were historically considered to be the simplest phenomenon from which to study epilepsy[9], and have been shown to perturb ongoing cognitive processes[10]. Therefore, IEDs represent a promising candidate to better understand the synchrony of the epileptic brain through its activity during small perturbations. Quantifying consistent changes in synchrony during IEDs would represent an important step towards understanding the underlying properties of the epileptic brain that accompany seizures.

In EEG recordings, IEDs are single sharp, spike-like waveforms that result from a burst of firing in a small group of neurons[8]. IEDs have a quantifiable impact on neural dynamics and behavior. For example, these focal perturbations affect a battery of cognitive tasks, even when those IEDs occur outside the tissue that supports seizure formation, typically referred to as the seizure onset zone (SOZ)[10, 11]. Additionally, IEDs are associated with spindle[12] and neural spiking[13] activity in regions distant from the source of the IED. These findings suggest that IEDs impact dynamics outside the population of cells producing the spike; however, a direct quantification of that impact in a large sample of source-level recordings has proven difficult, in part due to the scarcity of adequate data. Work using multiple imaging modalities has indirectly addressed this question by trying to separate out the contribution of IEDs to group differences in synchrony[14, 15] observed between individuals with epilepsy and controls. Some work suggests that differences in intrinsic functional connectivity between individuals with and without epilepsy are largely due to IEDs, rather than to endogenous dynamics[16]. However, results vary across cohorts and methods for quantifying synchrony[17]. Many studies have also detailed local or system-level changes in synchrony associated with IEDs, especially spatially near the onset of IEDs[18–22]. These findings are corroborated by biophysical modeling studies showing a lack of global differences in functional connectivity associated with IEDs, but some local changes[23]. However, quantifying only local changes makes it difficult to connect these findings to the global synchrony seen in diverse seizure states.

Studies of selective measures of intracranial EEG (iEEG) connectivity connectivity show that networks from IEDs activity tend to be more spatially similar to networks from seizure activity than seizure-free activity[24]. Suggesting that IEDs do change global network configuration, that those changes look more like seizure networks. A thorough investigation of the effect of IEDs on global synchrony is warranted to evaluate the response of an epileptic brain to perturbations. These observations could illuminate principles underlying the universal hypersynchrony seen across diverse seizure mechanisms. Here, we consider a thorough investigation to have two important properties: (1) it would comprehensively test different quantifications of global synchrony and (2) it would use a large sample to identify changes that are consistent across individuals. A comprehensive test would nearly exhaustively explore all relevant quantifications and aspects of global synchrony. Additionally, it would explicitly the results of all tests, regardless of if differences in synchrony are found or not. An investigation that meets these criteria would then serve as a thorough methodological resource to the research community who had questions about specific analyses, while also adding to our understanding of how synchrony changes during IEDs. But how will we quantify global synchrony?

Tools to quantify the response of a system such as the brain will often separately assess the effects of perturbations on different time scales of the system’s natural dynamics[25]. When the system is known, the prominent time scales can be extracted exactly as different periodic functions[26]. Here, since we only have access to the system’s outputs, we instead investigate time scales common to neural analyses through Fourier methods. In EEG recordings, synchrony at all of these time scales is quantified by an array of functional connectivity measures that identify statistical similarities between two signals. Additionally, EEG recordings provide information about synchrony in low frequency, high frequency, or broadband activity, each of which can reflect distinct neural processes. Determining precisely which processes are involved in each frequency requires a careful parameterization of both the frequency- and time-domain properties of a signal[27]. Here, we acknowledge that changes in synchrony likely arise from multiple neurophysiological processes, and therefore seek to discuss potential roles from across different neural mechanisms of each portion of the spectrum. Low-frequency activity (theta, alpha, beta) in a single region is thought to reflect aligned fluctuations in the membrane potential of a large, local population of neurons. When these bands show true oscillations, these fluctuations can modulate spiking activity to occur at the peaks of oscillatory rhythms[28]. Distant regions with similar low-frequency activity, and therefore high functional connectivity, could be structurally connected regions with high-amplitude oscillations where local spiking may or may not be modified[29]. Some theories of cortical communication suggest a role for these spatially broad oscillations in top-down processing and attention[30, 31]. Higher frequencies (especially high-gamma) in a single region are thought to correlate, though imperfectly[32], with local spiking activity[33, 34]. Multiple regions showing similar high frequency activity, and therefore high functional connectivity, are thought to have similar local spiking patterns. Both gamma and high-gamma connectivity are theorized to support bottom-up processing[30, 31]. Identifying which, if any of these processes are changing during IEDs could help connect basic principles of hypersynchrony to underlying biological processes that generate activity.

Here, we investigate 145 individuals from a publicly available dataset of individuals with epilepsy and iEEG recordings, to quantify changes in functional connectivity associated with IEDs that are not explained by changes in regional activity (**Fig. 1A**). We quantify connectivity based on multiple measures and multiple frequency bands (**Fig. 1B**). We use linear regression to quantify the amount that each of these metrics changes based on the presence of an IED, the number of IED sequences in a time window, and the average number of contacts containing an IED in each window (**Fig. 1CD**). This information allows us to determine whether there are consistent changes to functional connectivity associated with IEDs, whether these effects are band or measure specific. After identifying increases in functional connectivity associated with IEDs, we ask whether these changes are driven by specific edges. We first hypothesize that increases might be driven by changes to the tails of the distribution that capture the weakest and strongest edges. Given that interictal connectivity is characterized by a globally disconnected[35], but strongly intra-connected[36] seizure onset zone, we further hypothesized that increases might be driven selectively by changes within the seizure onset zone. After testing these hypotheses, we explore patterns of similarity across measures and bands. We conclude by assessing possible sources of individual differences in the magnitude of IED effects. This work provides a thorough quantification of changes in global functional connectivity beyond changes in activity associated with different properties of IEDs, and identifies several candidate neural processes that are disrupted during these perturbations. These observations contextualize IEDs within the complex ongoing dynamics of the brain, and can be used to validate models that test principles underlying the hypersynchrony in epilepsy.

**FIG. 1.**
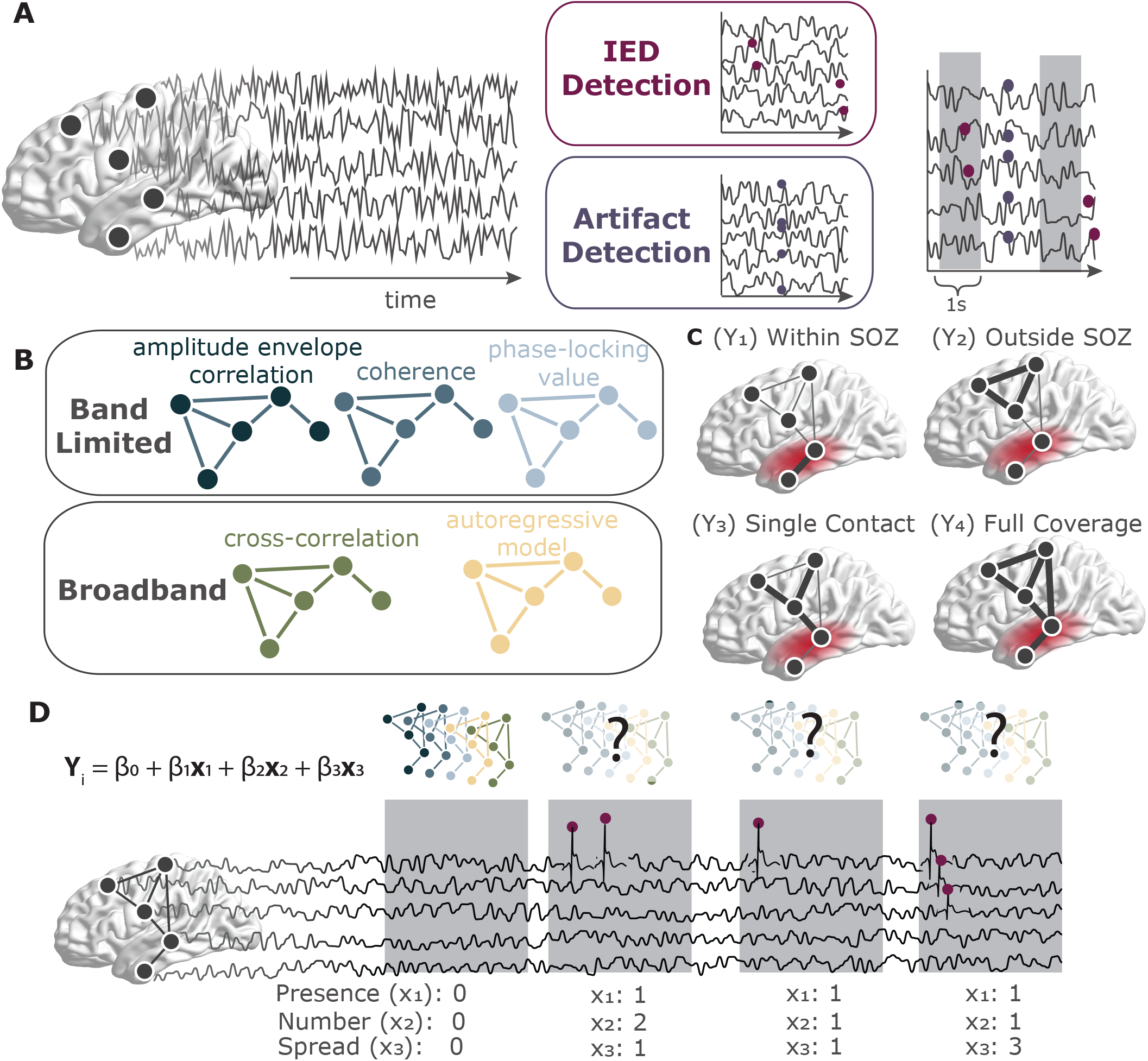
Schematic of methods. (**A**) Data undergo automatic IED and artifact detection before being separated into 1-second windows. (**B**) Description of the different functional connectivity measures being tested. (**C**) Description of the different summaries of global functional connectivity being tested. Red region indicates the seizure onset zone. (**D**) Schematic of regression equation used to estimate effect sizes for three different properties of IEDs: their presence, number, and spread.

## METHODS

### Participants (RAM dataset)

A publicly available dataset of 251 individuals undergoing intraoperative monitoring of their focal seizures was obtained from http://memory.psych.upenn.edu/RAM. In each participant, monitoring was conducted with either grid electrocorticography electrodes, stereoelectroencephalography electrodes, or both. Recordings were obtained while participants completed one of five different tasks testing either memory or free recall. Some participants underwent stimulation during these tasks. This data was collected as part of a multi-site, collaborative effort at the University of Pennsylvania, Columbia University, Dartmouth College, Emory University, Thomas Jefferson University, the Mayo Clinic, the National Institutes of Health, the University of Texas Southwestern, Medtronic Inc., and the Lawrence Livermore National Labs. A flow chart of preprocessing steps is shown in **Fig. S1**.

#### Individual-Level variables

We investigated the relative efficacy of twelve variables in explaining changes in functional connectivity associated with IEDs: age, sex, handedness, race, locus of IEDs, hemisphere of IEDs, etiology, the presence of a lesion, the age of seizure onset, the participants’s average rate of IEDs, the institution of treatment, and the type of contacts (grid or depth electrodes). Of these variables, sex, race, institution, age, type of coverage, and handedness were included as they were recorded in the public data release. If participants had missing values in any of these demographic variables, they were filled in with the most common demographic for categorical variables, or the mean demographic for continuous variables. The participant’s average rate of IEDs was calculated by dividing the average number of IEDs per window by the number of windows. Other variables used in later analyses required processing before including them in the model or were not included in the public data release. These will be described below.

We received additional clinical information that was not part of the public release from the creators of the original study. These data included: age at seizure onset, presence of a lesion, and the underlying etiology. The underlying etiology was determined by the epileptologists of the participating institution. Categorized used are as follows: (1) traumatic brain injury (*n*=53); (2) infection (*n*=18; *e.g*., herpes simplex virus, bacterial abscess); (3) neurocutaneous syndrome (*n*=2; including tuberous sclerosis); (4) neoplasia (*n*=4; e.g., tumor of glial origin); (5) stroke (*n*=3); (6) malformation of cortical development (*n*=35; including focal cortical dysplasia, heterotopia, polymicrogyria); (7) medial temporal sclerosis (*n*=6); (8) hypoxemic ischemic encephalopathy (*n*=6); (9) *other identified etiologies* in a different category (*n*=67), which included patients who may lack localizing lesions and/or did not accurately fall into the above-listed designations, such as those with epilepsy secondary to autoimmune (NMDA encephalitis), metabolic, vascular (developmental venous anomaly) or genetic causes; (10) unknown (*n*=128; without relevant abnormalities on examination, cognition, history or MR Imaging); or (11) multiple etiologies (*n*=17). The variable with missing values for the most people was etiology, which was missing in 79.3% of participants. Here, we note that these missing values are different from the unknown etiology, which notes that clinicians were unable to confidently determine etiology prior to iEEG investigations, rather than that information regarding etiology was not recorded. Individuals with missing values were not included in our final analysis.

In various analyses, we have used information about the locus, hemisphere, and tissue type (grey or white matter) for each contact. The locus and hemisphere of IEDs were obtained from information provided in the public data release. Coordinates provided in Montreal Neurological Institute (MNI) space were registered to the Schaefer parcellation[37] of seven cognitive systems[38]. It is standard practice to register atlases to a participant-specific space, rather than registering participant coordinates to a common space. To confirm the accuracy of system assignments, once each contact had been assigned a system, we manually checked that the physician-assigned regions (the ‘region’ field in the data release) of each contact matched their given system. Participants with missing MNI coordinates were excluded from these analyses (26.2%). White and grey matter labels for each contact were obtained from the Neuromorphometrics cortical parcellation protocol[39] given in the ‘wb’ field of the data release.

### Preprocessing electrocorticography data

First, raw data from the RAM dataset was segmented into 5-second or longer task-free epochs from either before or after task completion. If no information was available regarding the timing of task events, or if this information was inconsistent, the recording session was not processed. The median amount of data for each participant was 465.5 seconds r(ranged from 5.5 to 8,191 seconds). Data were then downsampled to the lowest sampling rate used across recording sites – 500 Hz – using the `resample()` function in MATLAB. Electric line noise and its harmonics at 60, 120, and 180 Hz (all sites were located within the United States) were filtered out using a zero phase distortion 4th order stop-band Butterworth filter with a 1 Hz width. This filtering was implemented using the `butter()` and `filtfilt()` functions in MATLAB. For impulse and step responses of this filter, see Supplemental **Fig. S2**.

We then sought to remove individual channels that exhibited excessive noise or had poor recording quality. The size of the dataset prevented us from visually inspecting each recording thoroughly; instead, we rejected channels using both the notes provided within the RAM dataset and via strict automated methods. After removing channels marked as low quality in the notes, we further removed electrodes that had either (1) a line length greater than three times the mean[10], (2) a *z*scored kurtosis greater than 1.5[40], or (3) a *z*-scored power-spectral density dissimilarity measure greater than 1.5[41]. The dissimilarity measure was the average of one minus the Spearman’s rank correlation of that signal with the signals of all other channels. These automated methods were chosen to remove channels with excessive high-frequency noise, electrode drift, and line noise, respectively.

Data were then demeaned and detrended. Detrending was used instead of a high-pass filter to avoid inducing filter artifacts[42]. Channels were then grouped by grid or depth electrode, and common average referenced within each group. Following the common average reference, plots of raw data and power spectral densities were visually inspected by an expert researcher with 7 years of experience working with electrocorticography data (J.S.) to ensure that data traces were of acceptable quality.

### Automatic interictal epileptic discharge (IED) detection

Automatic IED detection is still an open area of research, and there is as yet no consensus on best practices[43]. In clinical settings, epileptologists still manually mark EEG recordings for spikes and seizures when monitoring patients, although levels of agreement across different experts are not high [44]. We chose to use a previously reported IED detector from Janca *et al*.[45] because it is sensitive (89.3% sensitivity, 5.2 false positive rate), fast, and requires relatively little data per participant. This Hilbert-based method dynamically models background activity and detects outliers from that background. The initial *k* (threshold) value is set to 3.65, which was determined through cross-validation in prior work[45].

In order to remove false positives potentially caused by artifacts, we apply a spatial filter to identified IEDs. Specifically, we remove IEDs that are not present in a 50 ms window of IEDs in at least three other channels. The 50 ms window was taken from papers investigating the biophysical properties of chains of IEDs, which tended to last less than 50 ms[46]. Spikes detected within 50% of contacts within 2 ms were discarded. A subset of spikes were then randomly selected and validated by a board certified epileptologist (K.D.).

In order to quantify the spread and number of IEDs, we also determined where sequences of temporally proximal IEDs began and ended, using a previously utilized algorithm[7, 47]. IEDs occurring within 50 ms of the first IED in the sequence, or within 15 ms of the previous IED were considered part of the same sequence. We tested two additional biologically plausible parameter sets: IEDs occurring within 30 ms of the first IED or within 5 ms of the previous IED; and IEDs occurring within 100 ms of the first IED or within 30 ms of the previous IED. We found that both of these parameter sets resulted in identical definitions of IED sequences, demonstrating that our results are robust to different approaches used to operationalize a sequence. Sequences were discarded if 50% of the spikes in the sequence occurred within 2 ms of each other.

#### Alternate IED detector

To ensure that our results were not due to the specific IED detector used, we reproduced our key findings using an alternate detector. The alternate detector was the Delphos (Detector of ElectroPhysiological Oscillations and Spikes) detector[48]. This detector was designed to detect and distinguish between both oscillations and IEDs in the time-frequency representation of the signal[48]. Here, we chose a threshold value of fifty because it identified a similar number of IEDs as our primary detector. Hence, we knew that changes between detectors would not be driven by a different number of IEDs, but by differences in the properties of the individual IEDs discovered.

### Transient temporal artifact rejection

Visual inspection of the data revealed two types of temporally transient artifacts that we sought to remove from further analyses. The first type was sharp channel drift, which can be caused by physical movements of the participant, resulting in tugs of contacts and cables. Here, most channels simultaneously jump to a higher voltage before slowly drifting back to their original level. These sharp transients can cause artifacts that mimic oscillations when filtered[42]. To automatically detect these sharp artifacts, we calculated the rate of change of the timeseries and looked for large outliers. An absolute threshold of 30,000 microvolts per sample was used, as it was determined to be well outside the normal range for several randomly selected datasets, while still capturing artifacts (**Fig. S3A**). All time points where at least half of all channels contained values greater than this threshold were removed from further analyses.

The second type of artifact we observed was the presence of large periods of flatlining across all channels. These artifacts are less common and their origin is less certain, although they could be the result of a momentary disconnection between the amplifying and the recording systems. This flatlining could induce artificially large connectivity estimates as well as increased false positives in the IED detector. Since the detector dynamically calculates threshold values, periods of flat lining will lead to especially low thresholds and therefore many detected IEDs. Therefore, in addition to removing the flatlining periods themselves, we also removed 5 seconds of data following the artifact. To identify periods of flatlining, we searched for time points in the data with extremely small variance across all channels. We used a value of 300 microvolts because it was shown to be outside of the normal range, but able to capture flatlining for several randomly selected participants (**Fig. S3B**). All identified artifacts were visually inspected by an expert researcher with 7 years of experience working with electrocorticography data (J.S.) to ensure that identification was working as expected. After these steps, participants had a median of 416.8 seconds of data (ranging from 0 to 8,191 seconds).

### Dataset rejection

Our last quality control effort consisted of removing from further analysis entire datasets that were especially noisy. This step is important because if noise was present in the entire dataset, previously describes channel rejection methods relying on deviations from the mean would not remove all the sources of noise. Visual inspection of the data revealed three features indicative of low-quality data: (1) rough rather than smooth power spectral densities, (2) power spectral densities with many sources of line noise outside 60Hz and its harmonics, and (3) many temporal artifacts. To identify datasets with uncharacteristic power spectra, we first calculated the average power spectral density across all channels using Welch’s method in 500 ms windows (`pwelch()` in MATLAB). We then calculated the pairwise Spearman’s rank correlation coefficient between the power spectral densities of all datasets. We then flagged the 10% of datasets with the lowest correlations. The 10% threshold was chosen because it was the lowest threshold that removed all highly atypical power spectral densities (PSDs) upon visual inspection.

To remove datasets with high levels of line noise, we started with the same power spectral densities calculated above, and sought to identify those that were not fit well with a smooth curve. We then fit a smooth spline function to each power spectral density using `fit()` in MATLAB with a smoothing parameter of 0.01 Hz. Frequencies with notch filters and frequencies below 10 Hz were excluded from the fit, since they often deviated from the smooth curve and increased error rates. For each dataset, we then calculated the sum of squared errors between the smooth fit and the power spectral density. Power spectral densities with line noise at many frequencies would not be fit well by the smooth curve, and have higher errors. We then flagged the 10% of datasets with the highest error. Plots of the average retained power spectral densities of each dataset for windows with and without IEDs, and distributions of their corresponding aperiodic components can be found in **Fig. S4**.

Lastly, we flagged any dataset with greater than one-thousand time points containing temporal artifacts. Most of these datasets were also removed by one of the other methods. Lastly, we visually inspected the remaining data to confirm that the retained datasets looked clean. Any dataset that had been flagged during this step was then removed. After performing these steps, we were left with data from 181 participants.

### Dynamic functional connectivity

In this work, we seek to quantify how functional connectivity between brain regions changes in association with IEDs. To test for these changes, we first split all task-free data into 1 second non-overlapping windows. If a window contained an IED, that window was realigned to start one sample before the first IED in that window. This choice means that all IED windows will contain at least one IED sequence, and approximately 1 second of data following that IED. Successive windows would be shifted later, so that they did not overlap. Once windows were defined, data were prewhitened within each window using the Fieldtrip Toolbox for MATLAB[49] (`ft preprocessing()` function with the `derivative` parameter). Prewhitening reduces the autocorrelation of the timeseries, which in turn reduces the sampling variability of the connectivity estimates[50] To systematically explore the changes in functional connectivity following IEDs, we calculated five different commonly used metrics. For band-limited measures in the theta (*θ*, 4-8 Hz), alpha (*α*, 9-15 Hz), beta (*β*, 16-25 Hz), and gamma (*γ*, 36-70 Hz) bands, we wished to investigate amplitude-based, phase-based, and combined metrics (band definitions taken from Refs. [51, 52]). Therefore, we calculated orthogonal amplitude envelope correlations (amplitude), imaginary phase-locking value (phase), and imaginary multitaper coherence (combined). We also calculated the power in each band for each window, which we include as a covariate in later analyses. The power values can reflect both oscillatory power and changes in the aperiodic component of the signal. For simulations demonstrating that changes in the aperiodic component of the signal alone do not unduly impact our analyses, see Supplemental Analyses (**Fig. S5-S8**). For broadband measures, we sought to characterize both undirected functional connectivity and directed effective connectivity. Therefore, we calculated the maximal cross-correlation, as well as a vector autoregressive model.

#### Bandpass filter orthogonal amplitude envelope correlation

The amplitude envelope correlation (AEC) quantifies the extent to which signals from two channels change amplitude synchronously. For each bandpass filtered timeseries *y*_*n*_(*t*) at channel *n*, the instantaneous amplitude is obtained from the analytic signal *z*_*n*_(*t*) where 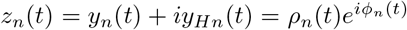. Here *φ*_*n*_(*t*) is the instantaneous phase and *ρ*_*n*_ is the narrow band waveform. Correlations between these amplitude envelopes are highly susceptible to artificial correlations due to volume conduction[53, 54]. In electrophysiological signals specifically, similarities between channels at location A and B could be caused by the signal from a third source C that spreads through the tissue and cerebrospinal fluid and is picked up by both channels, despite region C having no functional relationship to regions A and B. This process is called volume conduction, and while it is a much more relevant problem for sensor-level recordings (EEG and magnetoencephalography (MEG)), it can still have effects on iEEG data. Therefore, we take only the orthogonal components of each signal before calculating their correlation.

Here, we account for volume conduction using the method from Nolte *et al*. [54]. We chose this method as opposed to the method from Hipp *et al*. [53], because it uses a global normalization constant rather than one fit to each time point and is therefore much faster. Assuming Gaussian distributed data, which is a reasonable but not perfect fit to short segments of electrophysiological recordings, we can obtain the portion of the analytic signal *z*_*m*_ that is orthogonal to *z*_*n*_ by subtracting *z*_*n*_ multiplied by the real part of the coherency spectrum between the two channels. Specifically, we first normalize the analytic signal from channels *n* and *m* such that *<* |*z*_*n*_|^2^ *>*=*<* |*z*_*m*_| ^2^ *>*= 1. Here, the expected value is taken over time points. We then calculate coherency *c* as 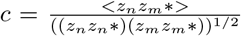 where refers to the complex conjugate. Lastly, we calculate the correlation between *z*_*n*_ and *z*_*m*_ − *real*(*c*)*z*_*n*_. The resulting value ranges from -1 to 1, with 1 indicating perfectly correlated signals, and -1 indicating perfectly anti-correlated signals. The absolute value of the orthogonal amplitude envelope correlation was taken before averaging across all pairwise estimates, or *edges*, in the functional network.

#### Bandpass filter imaginary phase-locking value (PLV)

The phase-locking value quantifies the temporal consistency of the phase offset between two channels regardless of the signal amplitude. The imaginary phase locking value removes 0-phase lag contributions to this value, which could arise from volume conduction. Using the bandpassed signal, *y*_*n*_(*t*), obtained with the parameters listed above, we calculate the imaginary phase locking value as 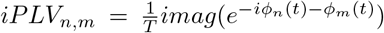. Here, *φ*_*n*_is the instantaneous phase in channel *n* for a given frequency band. This process was implemented using custom code in MATLAB, taken from Ref. [55]. Instantaneous phase requires a narrow frequency range in order to be biologically interpretable; therefore, we do not compute this measure on the high gamma band[56]. This measure ranges between 0 and 1, where 1 indicated consistent and small phase offsets, and 0 indicates inconsistent or large phase offsets.

It is possible for two signals to have meaningful phase synchrony in a given frequency range even if the power in the frequency range does not exceed the noise background (for neural signals, this background is the aperiodic component of a signal). However, the biological interpretation of such phase-locking is different, and less clear than phase locking in the presence of strong oscillations above the aperiodic background. Because of this difference in interpretation, we calculated the percentage of contacts that have oscillations above the aperiodic background in each window. We find that a minority of edges are between contacts with oscillations (1-9%), and that this number is variable across windows (**Fig. S9**).

#### Multitaper fourier transform – imaginary coherence & power

*Multitaper fourier transform:* Multitaper fast Fourier transforms (FFTs) use multiple tapers in order to better control spectral leakage at high frequencies[57]. They can be used to obtain the cross-spectral density, *S*_*n,m*_, between any two channels *n* and *m* as well as the power spectral density *S*_*nn*_ for channel *n*. The multitaper FFT was computed on thirty logarithmically spaced frequencies between 4 and 150 Hz using discrete prolate spheroidal sequences (DPSS) tapers with 4 Hz smoothing. Recording windows were zero-padded to the maximum period length for a given frequency. This calculation was implemented in MATLAB with the Fieldtrip toolbox[49].

#### Imaginary coherence

Coherence quantifies the consistency of phase offsets between two channels, weighted by their signal amplitude. Imaginary coherence ignores the contribution of 0-phase lag signals to this value, which could arise from volume conduction. Mathematically, the complex valued coherence *c* can be defined in multiple notations. Defined in terms of the cross-spectral density obtained via multitaper FFT, the coherence between two signals is 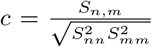 The imaginary coherence *C* is then taken as the absolute value of the imaginary component of coherence, *imag*(*c*). Imaginary coherence ranges from 0 to 1. Imaginary coherence values were averaged across all frequencies within a band.

##### Power

We wished to quantify changes to connectivity above what would also be explained by changes in activity. Here, we quantify activity as spectral power. Spectral power for each band and each channel was calculated by taking the logarithm (base 10) of all frequencies that fell within a given band.

#### Low-Pass filter – cross-correlation

##### Low Pass Filter

For broadband measures, we first removed sources of high-frequency noise using a low pass filter. Data were low pass filtered at 200 Hz using a 0-phase lag 4th order Butterworth filter. This filter was implemented using the MATLAB package Fieldtrip.

##### Cross Correlation

The cross-correlation is the maximal correlation between two signals that can be achieved across shifts of one signal relative to the other. The correlation coefficient was normalized such that the autocorrelation at zero lag was equal to one. The cross-correlation was implemented with the MATLAB function `xcorr()`. The resulting values range between 0 and 1.

#### Low-Pass filter – effective connectivity from autoregressive (AR) models

##### Autoregressive (AR) models

All the measures described above are undirected functional connectivity measures that will always provide the same estimates of the relation between node *n* and *m* as of the relation between node *m* and node *n*. We also wished to consider a directed measure of connectivity, in which the influence of node *n* on node *m* could be different from the influence of node *m* on node *n*. To accomplish this goal, we calculated a measure of directed connectivity from the weights of a first-order autoregressive model fit to the data. The weights were obtained by solving for (*A*) in the equation **x**(**t**) = **Ax**(*t* − 1) + *E*(*t*), where **x** is the timeseries data, and *E* is an error term. Fits were calculated using the arfit package in MATLAB[58]. Connectivity values from AR models can be positive or negative, so the absolute value was taken before summarizing. AR models assume a linear relationship between signals and assume the data are stationary. For short time windows (here, 1 second), this assumption is reasonable and has been made in previous work[59].

### Estimating effects of IEDs on functional connectivity

The overarching goal of this study is to quantify the effect of IEDs on each of the functional connectivity metrics defined above. We also wished to be able to answer the three specific questions: (1) Are changes in connectivity driven by shifts in the weakest or strongest connections? (2) Are changes larger within than outside the seizure onset zone? (3) Do effects of IEDs vary reliably across regions or individuals? To answer these questions, we calculated IED effects on five different summary measures of the distributions of connections. These measures included (1) the average connectivity across all contacts (full coverage), (2) the skew of connectivity across all contacts, (3) the strength of connectivity inside the seizure onset zone, (4) the strength of connectivity outside the seizure onset zone, and (5) the strength of connectivity in each individual contact (**Fig. 1C**). We note that, in every case, all contacts in the seizure onset zone are considered to be part of a single zone, even though it is possible that participants have multiple seizure foci. To estimate how many participants might have multiple foci, we calculated the distance between all contacts in the SOZ. We found that 5% (7) of individuals had multiple peaks, suggesting that multiple foci are not a large confounding factor in our analyses. In supplemental analyses, we also reported the changes in connectivity in two additional summary measures: (1-2) between all contacts inside and outside the irritative zone, (3-4) the strength between all grid and depth contacts separately, and (5-6) the strength between grey and white matter portions of depth contacts.

We next quantified the specific features of each IED sequence that we hypothesized would impact functional connectivity uniformly. The first feature was the presence of an IED in a window, regardless of the properties of that IED. The second was the number of sequences within a window. A sequence was defined as all IEDs occurring within 50 ms of the first IED, or within 15 ms of the last IED in that timeframe. If the leader channel had multiple spikes in this sequence, the initial sequence was split into multiple sequences. The third was the average spread across each sequence within the window (**Fig. S1D**).

For each dependent variable, we calculated the coefficient of each IED predictor in a permutation-based linear model that contained power and recording session as nuisance covariates. If two or more of the three covariates of interest were perfectly collinear, one of them was removed from the model. Typically, collinearity occurred when one or more predictors did not vary. For example if all windows contained only one IED, the presence and number of IED predictors would be collinear. The results presented in the main text are obtained from a subset of 145 participants that had coefficient values for all three predictors. Outliers, defined as values three standard deviations smaller or larger than the mean distribution for each band and measure combination, removed from further analyses. Permutation-based models were used because both the spike spread and the number of spikes are highly skewed, non-normal variables that could lead to artificially large estimations in parametric models. Additionally, permutation-based models more effectively down-weight outliers in the distribution.

### Group LASSO

We used group LASSO regression to assess the relative importance of each included variable. Group LASSO applies a penalty to groups of variables and will regularize coefficients of uninformative groups to 0. Here, all levels of a given categorical variable were grouped together. This way, all levels of the etiology variable, for example, would be regularized together, rather than regularizing individual levels. Group LASSO was implemented using the gglasso package in R (https://github.com/emeryyi/gglasso).

### Statistical analyses

Here, we set out to complete an exploratory analysis of the changes to functional connectivity associated with IEDs. We used a strict family-wise Bonferroni correction in which each band/measure combination (*n*=16) was treated as a different hypothesis. The Bonferroni correction sets a new threshold for significance *α*_*corrected*_ for a set of *n* comparisons equal to *α*/n. Here, *α* is equal to 0.05. This procedure allowed us to have more confidence in the results we report, but its stringency may contribute to false negatives.

Before any significance testing, participants with changes to functional connectivity greater than three standard deviations above the mean were removed. Here, the phrase *changes in functional connectivity* refers to the coefficient from the regression of IED features (e.g., presence, number, and spread) from strength. Quantile-quantile plots were then used to check whether distributions were normal. Since distributions appeared to deviate from a normal distribution, one sample permutation tests assessed whether distribution means were significantly different from zero (**Figs. 2-3**). One sample permutation tests were performed with the EnvStats package for R (https://github.com/alexkowa/EnvStats). The test consists of comparing the sign difference in the mean from zero to all possible permutations of signs. Statistics are reported with the test statistic, the number of observations, and the *p*-value. Similarity was assessed with Spearman’s rank correlations (**Figs. 4-5**) to mitigate the influence of outliers.

**FIG. 2.**
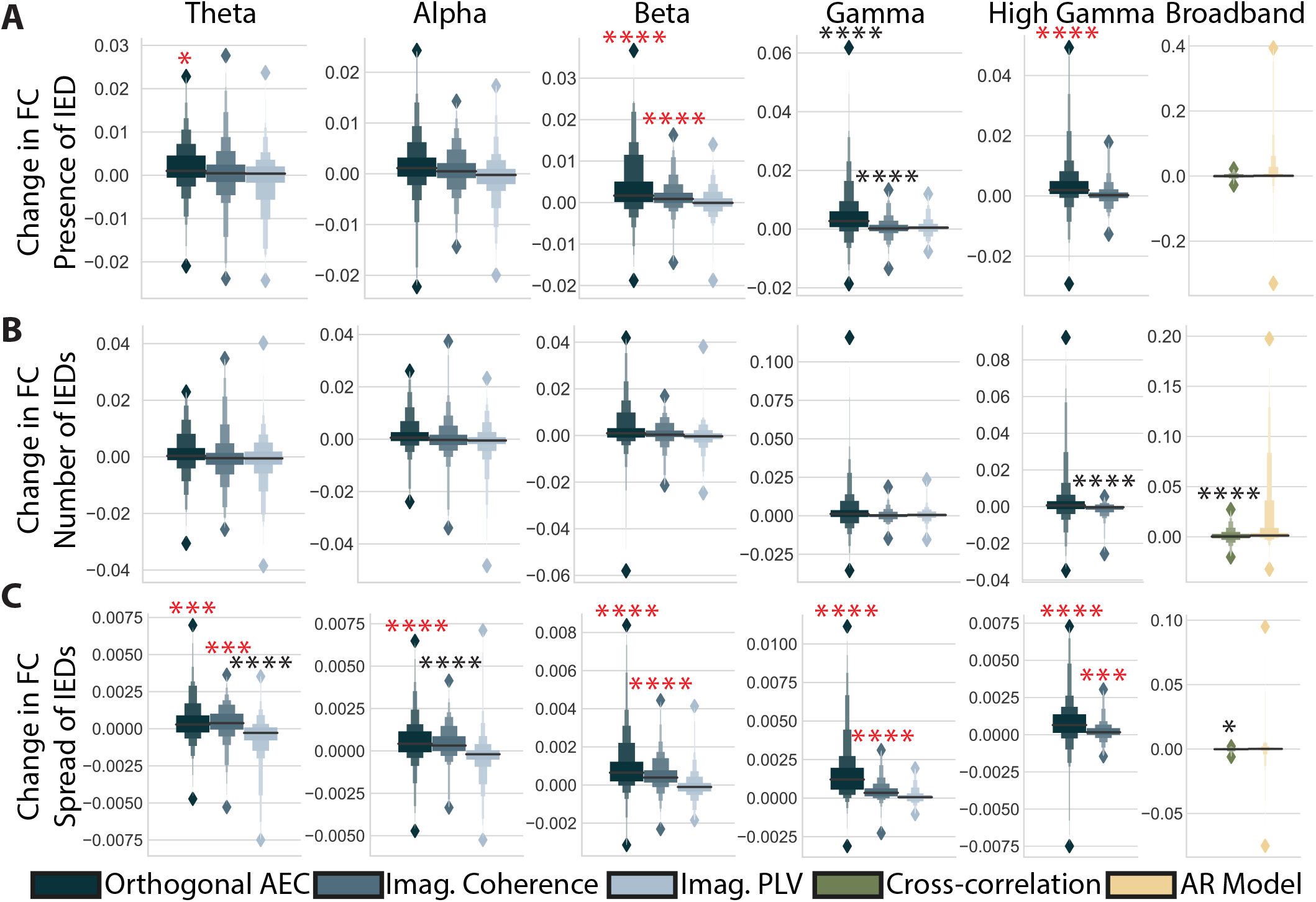
Changes in Functional Connectivity Associated with IEDs. (**A**) Magnitude of the change in functional connectivity (FC) for the presence of an IED obtained from permutation based regression including all IED predictors, power in a given band, and the recording session. Columns indicate different frequency bands, and colors indicate different measures. Bright red asterisks indicate significant distributions after multiple comparisons correction that could also be reproduced with a different spike detector. Black asterisks indicate effects that were not reproducible with another IED detector. * = *p <* 0.05, ** = *p <* 0.01, *** = *p <* 0.001, **** = *p <* 0.0001 (after Bonferroni correction (n=16). (**B**,**C**) Same as in panel *(A)*, but for the coefficients associated with the number of IEDs and the spread of IEDs. Here, AEC stands for amplitude envelope correlation, imag. for imaginary, PLV for phase locking value, and AR for autoregressive model.

**FIG. 3.**
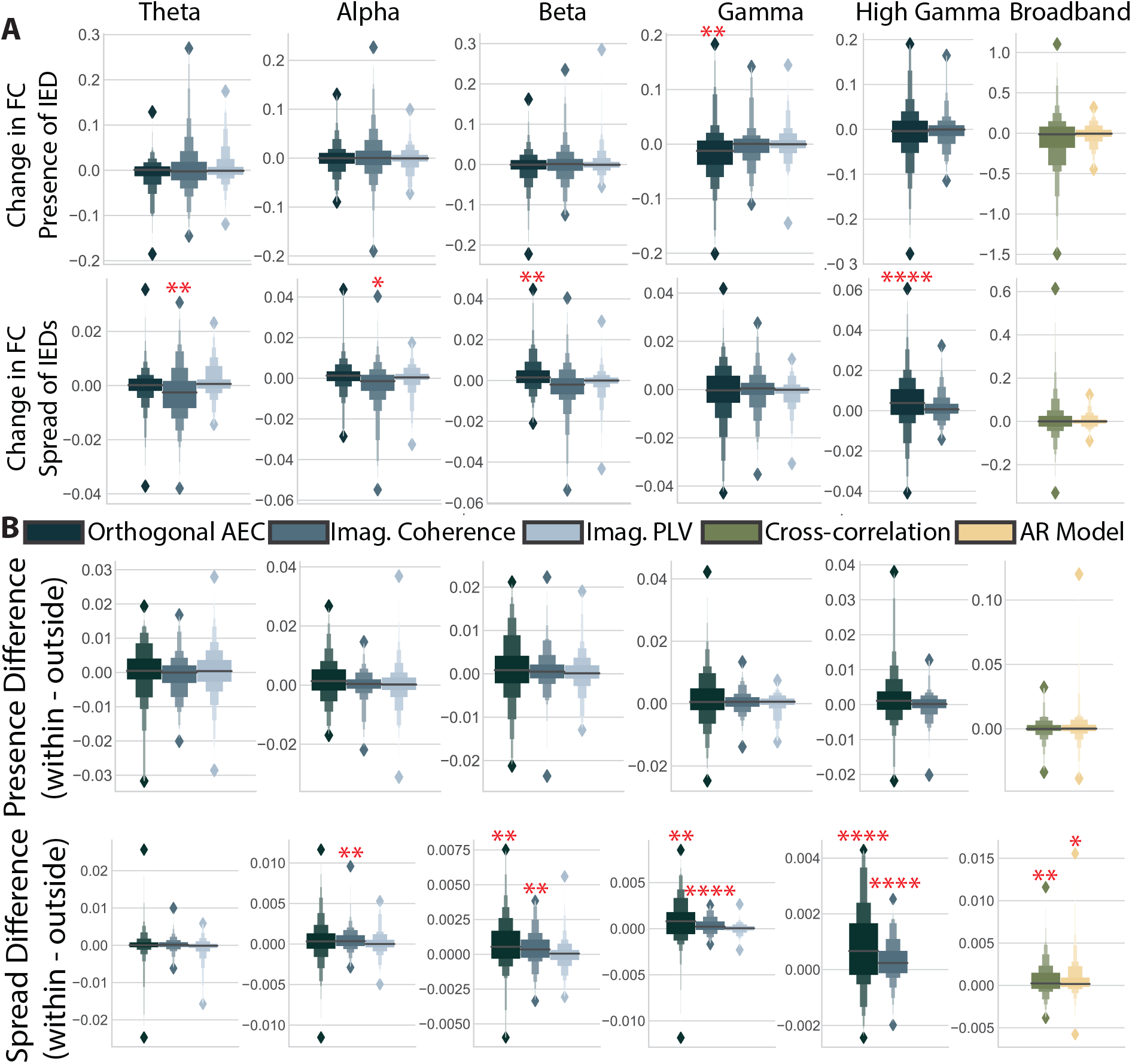
Changes to specific connections. (**A**) The coefficient associated with each predictor for explaining changes to the skew of the distribution of functional connectivity. * = *p <* 0.05, ** = *p <* 0.01, *** = *p <* 0.001, **** = *p <* 0.0001. Grey asterisks indicate significant differences before multiple comparisons correction (after Bonferroni correction (n=16)) (**B**) The difference between coefficients found for changes to connections within the seizure onset zone versus outside it.

**FIG. 4.**
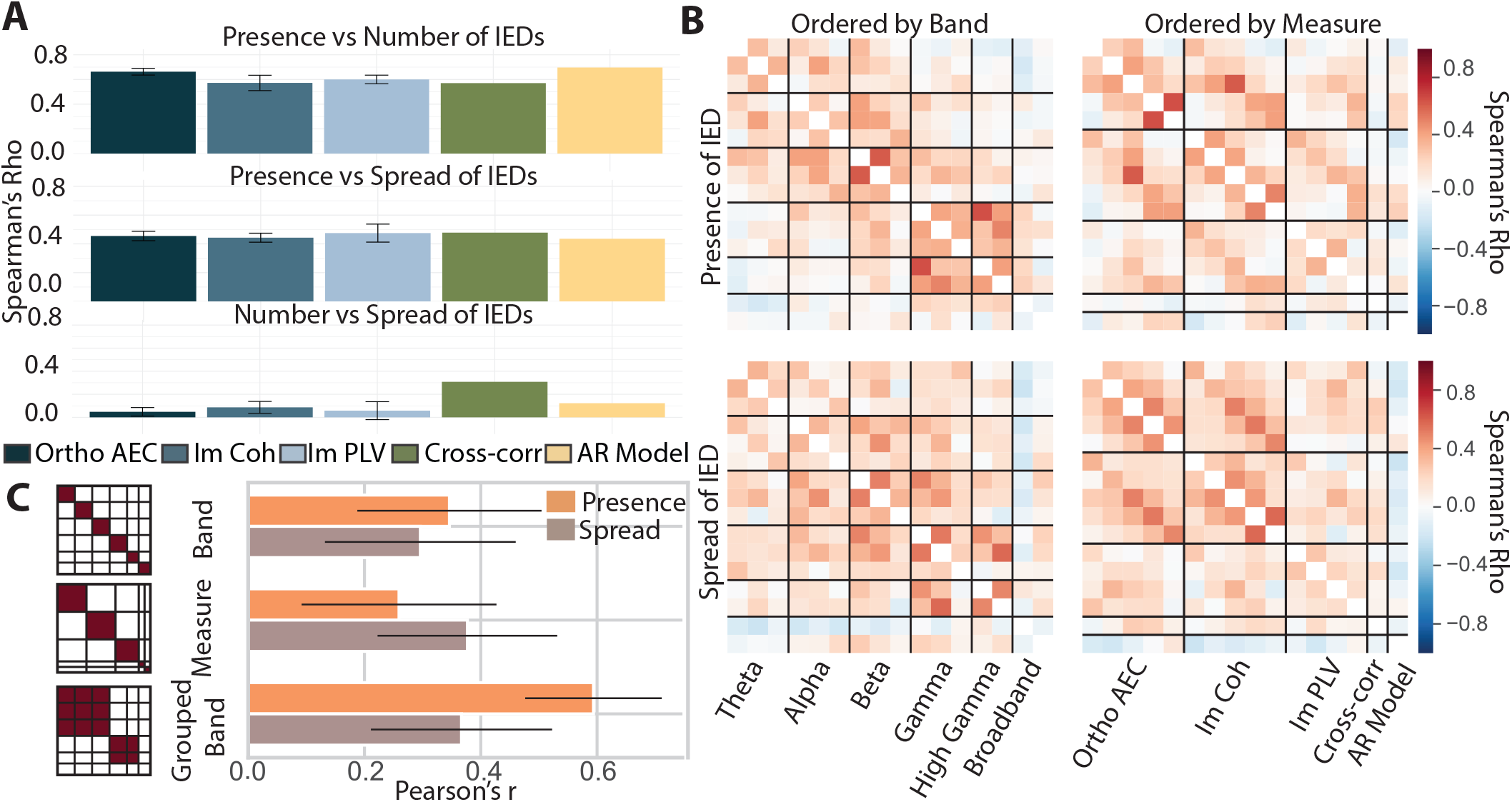
Similarity across predictors, measures, and bands. (**A**) Spearman’s correlations between predictors. A bar plot showing the average correlation and standard error for each measure across bands. Error bars are the standard error across bands, when there are multiple bands to test. Colors indicate different measures. (**B**) Similarity matrices between all band-measure combinations. *(Left)* Matrices ordered by band. *(Right)* Matrices ordered by measure. (**C**) Correlations with a frequency band mask *(left)*, measure mask *(middle)*, and frequency band group, i.e. high or low *(right)* for each predictor. Error bars show the 95% confidence interval for each correlation.

**FIG. 5.**
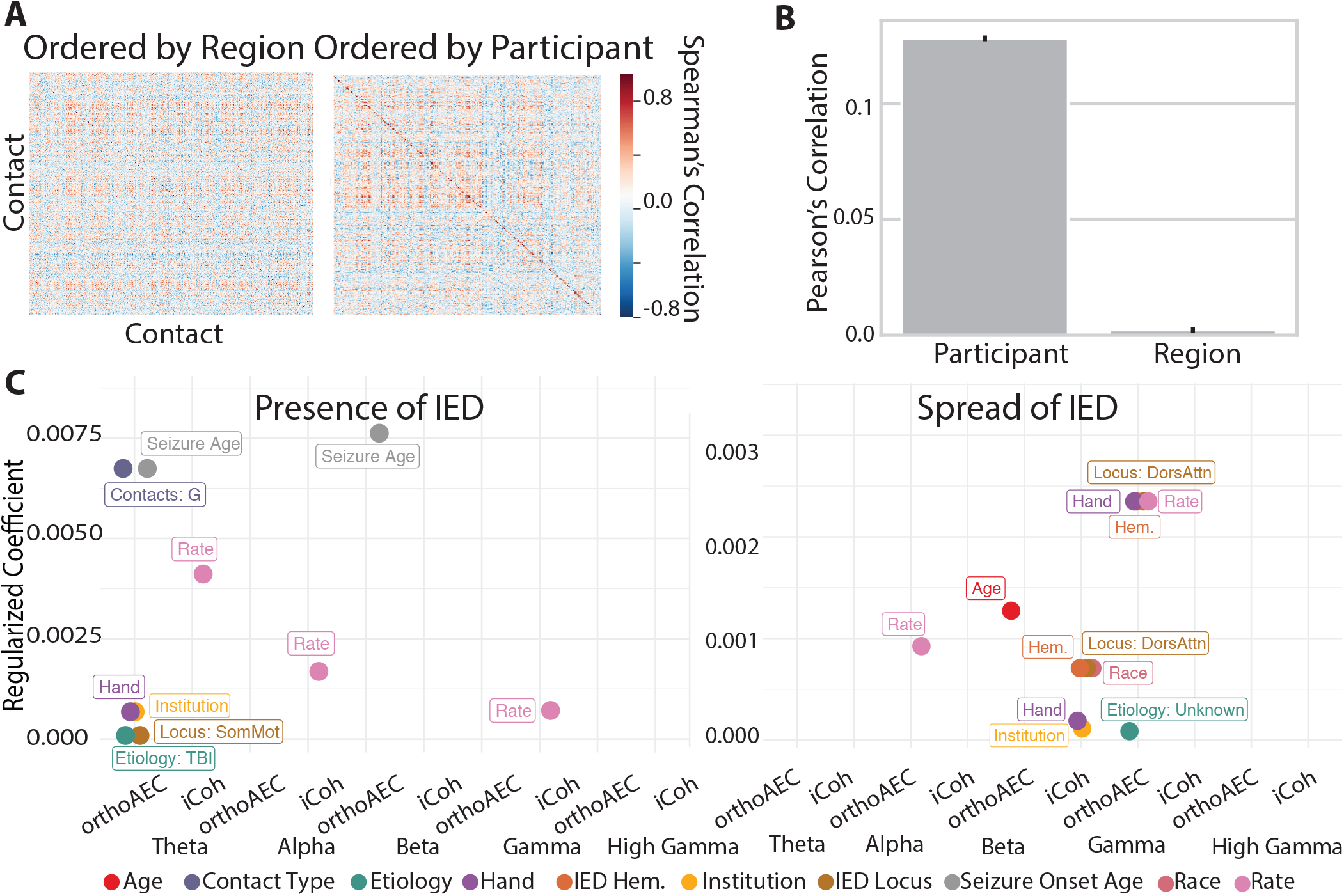
Individual differences. (**A**) Similarity matrices for coefficient distributions across all predictors, bands, and measures, but between contacts. *(Left)* Matrices ordered by region. *(Right)* Matrices ordered by participant. (**B**) Correlations for masks selecting for participants, or regions. Error bars are the 95% confidence interval for each correlation. (**C**) Predictors included for each model, and their corresponding regularized coefficient. For categorical variables representing clinical, or contact level variables, text is displayed showing the level of the categorical variable that had the largest coefficient. Different variables are shown in different colors, with a legend matching colors to variables at the bottom of the plot. Only included variables are shown in the legend; therefore, no color is assigned to the sex, or an indicator of whether of not each participant was lesional. Legend and text abbreviations are as follows: Hem. is an abbreviation for hemisphere, Contacts *G* indicates grid coverage; Etiology *TBI* indicates traumatic brain injury *SomMot* indicates the somatomotor network, and *DorsAttn* indicates the dorsal attention network.

Differences within versus between groups in similarity matrices were assessed with Pearson’s correlations between the lower diagonal of matrix entries, and a binary mask specific to each hypothesis in **Figs. 4-5**. This test is similar to those used in representational similarity analysis and are designed to test hypothesis about higher-order, rather than pairwise similarity structure[60]. To assess the significance of these measurements, we tested correlations calculated from empirical similarity matrices against correlations calculated from one-hundred permuted null models. In each null model, a random mapping of true band-measure pairings to permuted band-measure pairing was given to all participants.

### Data and Code

Code is available at github.com/jastiso/interictal fc. Data is publicly available at http://memory.psych.upenn.edu/RAM.

## RESULTS

We sought to characterize the changes in global functional connectivity associated with a simple form of epileptic activity – an IED – in individuals with medically refractory epilepsy. Accordingly, we used a large sample of 145 participants in the RAM dataset. We select a comprehensive sample of functional connectivity measures common to iEEG analysis. Specifically, we calculate three measures of band-limited functional connectivity and two measures of broadband functional connectivity in 1 second windows. Windows containing IEDs were aligned to start 1 sample before the IED sequence. Next, we quantified the effect of three different properties of IEDs on each band-measure combination, using a permutation-based linear model including the presence of an IED, the number of IEDs in a window, and the average spread of every IED in the window (**Fig. 1D**). In order to identify drivers of spatial distributions of observed effects, we calculated the effects of IEDs on the total strength across all contacts, the skew of edges across all contacts, the strength of only those contacts within (or outside) the seizure onset zone, and in each contact individually (**Fig. 1C**).

### Quantifying the impact of IEDs on functional connectivity

We first asked which band-measure combinations would show consistent changes in the strength of functional connections for each of the three IED predictors. We find different patterns of responses across predictors. The presence of IEDs increases functional connectivity, but largely in the orthogonal amplitude envelope correlations (**Fig. 2A**; Table T1. The number of IEDs in a window does not consistently affect functional connectivity. The only exceptions were the orthogonal amplitude envelope correlation in the beta band and high gamma band (**Fig. 2B**; Table T1). The spread of IEDs within a window largely increases both amplitude-based (orthogonal amplitude envelope correlation) and amplitude weighted phase-based (imaginary coherence) measures (**Fig. 2C**; Table T1). The predictor for the spread of IEDs also shows changes to functional connectivity that are an order of magnitude smaller than the other two predictors. Our results, based on a large sample (*n* = 145) of source-level data, demonstrate consistent increases in global functional connectivity as a result of the presence and spread of IEDs.

To ensure that results are driven by IEDs themselves, and not by artifacts due to our specific IED detection algorithm, we repeated the above analysis with a two sets of parameters in a second IED detector. This detector used spectrotemporal features rather than the magnitude of the bandpassed signal to identify IEDs. We repeated our analysis using one parameter selection that gave a similar number of IEDs to our original detector and one parameter selection that was recommended by the alternative detector. With these alternative approaches, we found significant effects for all but 7 of the measures listed above: the gamma amplitude envelope correlation and imaginary coherence (number); high gamma imaginary coherence (number); broadband cross-correlation (number); theta imaginary phase-locking value (spread); alpha imaginary coherence (spread); and broadband crosscorrelation (spread) (**Fig. 2**, significance indicated with dark red asterisks). Using this algorithm, we also identified four additional, inconsistent measures that had a mean significantly different from zero. These measures were the alpha orthogonal amplitude envelope correlation (presence), gamma imaginary phase-locking value (presence), gamma orthogonal envelope correlation (number), and the gamma phase-locking value (spread). This observation indicated that while the selection of IED detector can influence the quantification of neurophysiological changes associated with IEDs, the finding that functional connectivity increases during IEDs is consistent across detectors.

### Quantification in subsets of connections

We next asked if these global effects were driven largely by smaller parts of the network. Because we found no reproducible large changes to functional connectivity for the number-of-IEDs predictor, we limited this analysis to the presence and spread of IEDs (for the number of IEDs, see **Fig. S10**). We hypothesized that increases in global connectivity were driven by a strengthening of the weakest connections, possibly connecting the seizure onset zone to the rest of the brain, or by a strengthening of already strong, isolated connections within the seizure onset zone. We broke these hypotheses first into two questions: (1) could increases be driven by the weakest edges? And (2) could increases be driven by edges in the seizure onset zone? To address our first question, we repeated the above analysis with the skew, rather than the strength of edges. Since edge distributions are heavy tailed (see **Fig. S11**), increases in the skew are consistent with a strengthening of the weakest edges in the distribution. We observe far fewer significant changes to the skew of connections than to the strength, and we note that these changes tend to be negative. Significant changes were seen in gamma orthogonal amplitude envelope correlation (presence), broadband cross-correlation (number), theta and alpha imaginary coherence (spread), and beta and high gamma orthogonal amplitude envelope correlation (spread) (**Fig. 3A**; Table T2). These findings indicate that most changes observed above are not driven by a strengthening of weak connections.

**TABLE I.**
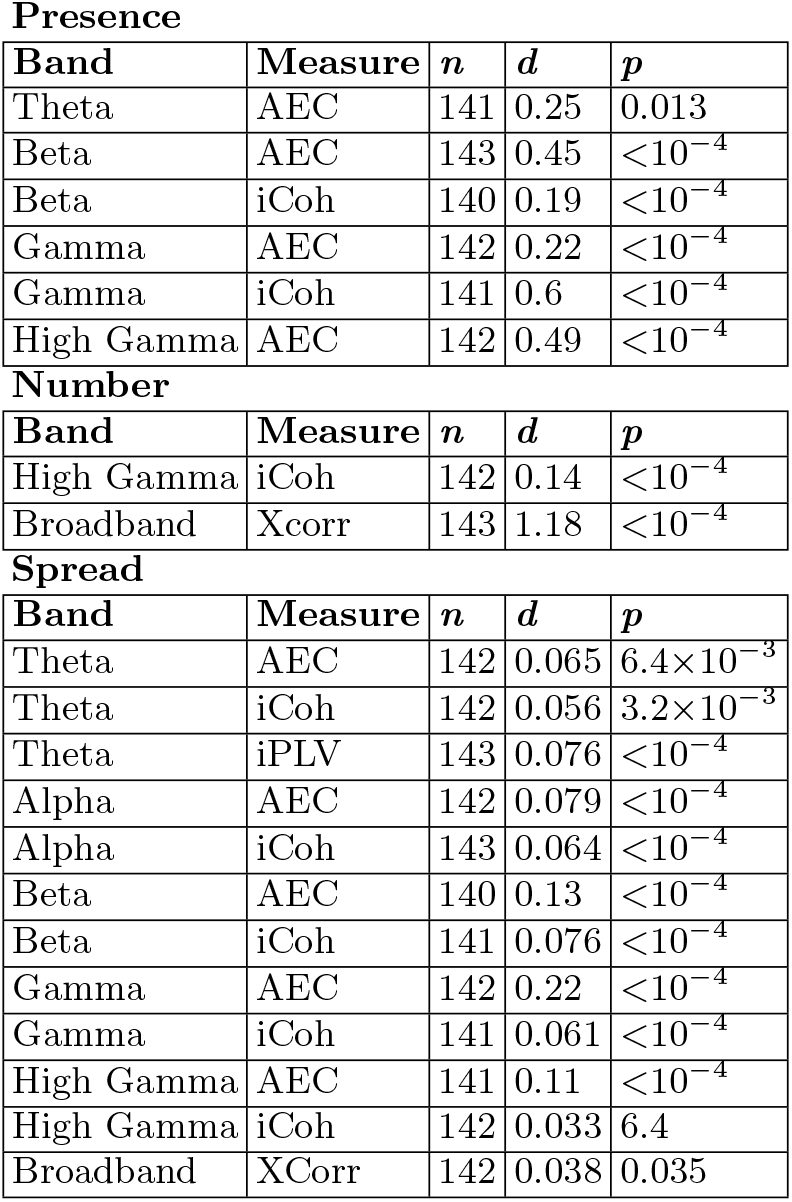
Statistics for changes in functional connectivity associated with IEDs. *n* is the number of observations. *d* is the effect size for the one-sized permutation test. *p* is the *p*-value. All *p*-values are Bonferroni corrected across the 16 repeated tests.

Presence
| Band | Measure | $n$ | $d$ | $p$ |
| --- | --- | --- | --- | --- |
| Theta | AEC | 141 | 0.25 | 0.013 |
| Beta | AEC | 143 | 0.45 | $<10^{-4}$ |
| Beta | iCoh | 140 | 0.19 | $<10^{-4}$ |
| Gamma | AEC | 142 | 0.22 | $<10^{-4}$ |
| Gamma | iCoh | 141 | 0.6 | $<10^{-4}$ |
| High Gamma | AEC | 142 | 0.49 | $<10^{-4}$ |

Number
| Band | Measure | $n$ | $d$ | $p$ |
| --- | --- | --- | --- | --- |
| High Gamma | iCoh | 142 | 0.14 | $<10^{-4}$ |
| Broadband | Xcorr | 143 | 1.18 | $<10^{-4}$ |

TABLE I. **Statistics for changes in functional connectivity associated with IEDs.** $n$ is the number of observations. $d$ is the effect size for the one-sided permutation test. $p$ is the $p$ -value. All $p$ -values are Bonferroni corrected across the 16 repeated tests.
| Band | Measure | $n$ | $d$ | $p$ |
| --- | --- | --- | --- | --- |
| Theta | AEC | 142 | 0.065 | $6.4 \times 10^{-3}$ |
| Theta | iCoh | 142 | 0.056 | $3.2 \times 10^{-3}$ |
| Theta | iPLV | 143 | 0.076 | $<10^{-4}$ |
| Alpha | AEC | 142 | 0.079 | $<10^{-4}$ |
| Alpha | iCoh | 143 | 0.064 | $<10^{-4}$ |
| Beta | AEC | 140 | 0.13 | $<10^{-4}$ |
| Beta | iCoh | 141 | 0.076 | $<10^{-4}$ |
| Gamma | AEC | 142 | 0.22 | $<10^{-4}$ |
| Gamma | iCoh | 141 | 0.061 | $<10^{-4}$ |
| High Gamma | AEC | 141 | 0.11 | $<10^{-4}$ |
| High Gamma | iCoh | 142 | 0.033 | 6.4 |
| Broadband | XCorr | 142 | 0.038 | 0.035 |

We address our second question by subtracting coefficients for the effect of IEDs on connections inside the clinically defined seizure onset zone (SOZ) from connections outside the SOZ, for participants where this data was available (*n* = 104). Large positive numbers would indicate greater changes inside the SOZ. We find that, after correction for multiple comparisons, only the changes in connectivity resulting from the spread of IEDs were larger in the seizure onset zone compared to the rest of the brain (**Fig. 3B**). While these differences are seen in all but the theta band, they are most pronounced in the connectivity of the high gamma band (**Fig. 3B**; Table T3). Some measures, indicated with a grey asterisk in **Fig. 3**, showed significant uncorrected differences, and also suggested larger effects inside the seizure onset zone. To determine whether the changes in global functional connectivity associated with the spread of IEDs was driven by edges within the SOZ, we tested whether edges between contacts outside the SOZ showed changes in functional connectivity that were statistically different from 0. Only one measure of the original twelve significant measures, the high gamma imaginary coherence, was no longer significant when only considering edges outside the SOZ. Overall, we find that only changes associated with the spread, and not the presence, of IEDs are increased within the seizure onset zone. While changes are larger within the SOZ, edges outside the SOZ still show positive changes associated with the spread of IEDs.

We then tested our third hypothesis: that changes would be driven by any contact that contained an IED rather than the seizure onset zone (SOZ). Changes in these contacts activity could arising from statistical properties the spikes themselves. We first calculated differences in connectivity between contacts in with and without IEDs. We then use the same methods as above to assess if this measure of strength changes with each IED preoperty. Positive values indicate higher connectivity between contacts with IEDs, and negative values indicate higher connectivity in contacts iwthout IEDs. We find five significant changes instrength after multiple comparisons correction for the spread of IEDs spanning all bands and measures (**Fig. S12**). In all but one case (theta iPLV) changes are negative, indicating greater connectivity outside of contacts with IEDs. Overall, these finding suggests that the increases in connectivity observed above are usually driven by contacts without IEDs.

**TABLE II.**
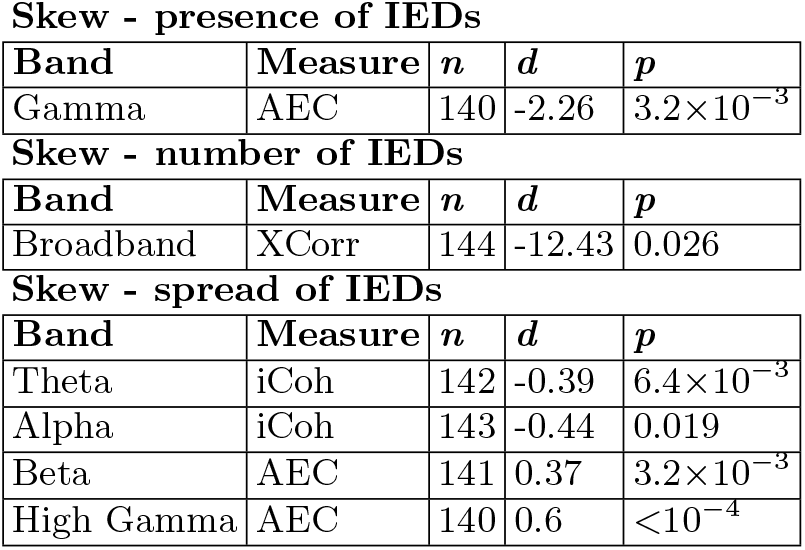
Statistics for changes in skew. *n* is the number of observations. *d* is the effect size for the one-sized permutation test. *p* is the *p*-value. All *p*-values are Bonferroni corrected across the 16 repeated tests.

**Skew - presence of IEDs**
| Band | Measure | $n$ | $d$ | $p$ |
| --- | --- | --- | --- | --- |
| Gamma | AEC | 140 | -2.26 | $3.2 \times 10^{-3}$ |

**Skew - number of IEDs**
| Band | Measure | $n$ | $d$ | $p$ |
| --- | --- | --- | --- | --- |
| Broadband | XCorr | 144 | -12.43 | 0.026 |

TABLE II. **Statistics for changes in skew.** $n$ is the number of observations. $d$ is the effect size for the one-sided permutation test. $p$ is the $p$ -value. All $p$ -values are Bonferroni corrected across the 16 repeated tests.
| Band | Measure | $n$ | $d$ | $p$ |
| --- | --- | --- | --- | --- |
| Theta | iCoh | 142 | -0.39 | $6.4 \times 10^{-3}$ |
| Alpha | iCoh | 143 | -0.44 | 0.019 |
| Beta | AEC | 141 | 0.37 | $3.2 \times 10^{-3}$ |
| High Gamma | AEC | 140 | 0.6 | $< 10^{-4}$ |

**TABLE III.** Statistics for differences within and outside the SOZ. *n* is the number of observations. *d* is the effect size for the one-sized permutation test. *p* is the *p*-value. All *p*-values are Bonferroni corrected across the 16 repeated tests.

| Band | Measure | $n$ | $d$ | $p$ |
| --- | --- | --- | --- | --- |
| Alpha | iCoh | 102 | 0.05 | $3.2 \times 10^{-3}$ |
| Beta | AEC | 101 | 0.07 | $3.2 \times 10^{-3}$ |
| Beta | iCoh | 102 | 0.043 | 0.019 |
| Gamma | AEC | 101 | 0.079 | 0.032 |
| Gamma | iCoh | 101 | 0.033 | $< 10^{-4}$ |
| High Gamma | AEC | 100 | 0.083 | $3.2 \times 10^{-3}$ |
| High Gamma | iCoh | 99 | 0.03 | 0.02 |
| Broadband | AR | 104 | 0.063 | $6.4 \times 10^{-3}$ |
| Broadband | XCorr | 101 | 0.066 | 0.03 |

Given that the changes in functional connectivity associated with the presence of IEDs had a similar magnitude within and outside of pathological tissue, we hypothesized that changes might be driven by edges that sample tissue differently, or that are located in different types of tissue. In particular, we hypothesized that changes in functional connectivity might be driven by connections amongst grid contacts versus depth contacts, or within grey versus white matter on depth contacts. For all individuals who had a combination of grid and depth coverage (*n* = 35), we assessed whether functional connectivity changes were greater for edges within contacts in the grid versus edges within depth contacts. Due to the smaller sample size, we only tested this effect in functional connectivity band and measure combinations that showed significant changes in functional connectivity (the orthogonal amplitude envelope correlation and imaginary coherence for all bands). We find no significant differences between edges within grid and depth coverage, either before or after multiple comparisons corrections (**Fig. S13**). We then repeated this process for individuals who had depth contacts marked as grey and white matter (*n* = 104). We found no significant differences between white and grey matter after multiple comparisons corrections, although some measures were significant before correction (**Fig. S13**).

### Relationship between IED predictors and changes to connectivity within frequency bands and connectivity measures

Earlier, we explored a large space of IED predictors and their effects on different frequency bands and connectivity measures. In order to distill these many findings into broader, unifying principles regarding the behavior of neural systems during simple epileptiform events, it would be useful to determine whether there are consistent associations between different predictors, frequency bands, or measures. To identify these associations, we calculate similarity with Spearman’s correlations of participants’ coefficients between IED predictors and band-measure combinations.

We first computed pairwise correlations between the distribution of participants’ coefficients for each of the IED predictors, in each band-measure combination. We find that relationships are largely consistent across bands, and we therefore show results from each measure averaged across bands (**Fig. 4A**). We find that the strongest effect is a significant positive correlation between the magnitude of changes associated with the number of IEDs and the presence of an IED (**Fig. 4A**; Spearman’s correlation, Bonferroni-corrected *p*-values (*n* = 16) mean *r*_*AEC*_ = 0.66± 0.061, mean *r*_*iCoh*_ = 0.57± 0.14, mean *r*_*iP LV*_ = 0.60 ± 0.070, *r*_*xcorr*_ = 0.57, *r*_*AR*_ = 0.70, all *p <* 5.14 ×10^−5^). The correlations between the connectivity changes associated with the presence of an IED and the spread of an IED tend to be significant and positive but weaker (**Fig. 4A**; Spearman’s correlation, Bonferroni-corrected *p*-values (*n* = 16) mean *r*_*AEC*_ = 0.45± 0.073, mean *r*_*iCoh*_ = 0.42 ± 0.070, mean *r*_*iP LV*_ = 0.47 ± 0.13, *r*_*xcorr*_ = 0.48, *r*_*AR*_ = 0.44, all *p <* 8.47 × 10^−3^), and the correlation between the number of IEDs and the spread of IEDs tends to be nonsignificant and much weaker (**Fig. 4A**; Spearman’s correlation, Bonferroni-corrected *p*-values (*n* = 16) mean *r*_*AEC*_ = 0.048 ± 0.083, mean *r*_*iCoh*_ = 0.086 ± 0.12, mean *r*_*iP LV*_ = 0.058 ± 0.16, *r*_*xcorr*_ = 0.31, *r*_*AR*_ = 0.12; only broadband cross-correlation and theta imaginary phase-locking value had signififcant *p*-values after multiple comparisons correction). Due to the lack of consistent changes in functional connectivity resulting from the number of IEDs and the predictor’s high similarity with the presence of an IED, we excluded this predictor from further analysis.

Next, we sought to characterize similarities between each band-measure combination. Here we test three hypotheses. First, we hypothesize that coefficients will be similar across measures within a given frequency band. Our second hypothesis is that coefficients will be similar within a measure, across frequency bands. Our third hypothesis is that there will be a patterned similarity across bands. Specifically, the low frequencies (theta, alpha, and beta) will be similar to each other, but dissimilar to high frequencies (gamma and high gamma) and *vice versa*. High and low frequencies are often described in the literature as showing opposing patterns, and are theoretically proposed to have opposite roles[30]. To test these three hypotheses, we calculate similarity matrices for each predictor. Each element of the similarity matrix is the Spearman’s correlation between the distribution of functional connectivity changes across participants for two band-measure combinations. We show these matrices sorted by frequency band and by measure (**Fig. 4B**).

To test our first hypothesis, we correlate the upper triangles of these pairs of similarity matrices with masks that operationalize each hypothesis. For example, in the band similarity hypothesis, the mask has ones for entries from the same band and zeros for entries from different bands. To test the significance of our findings, we compare the correlations calculated from empirical data to correlations calculated from one-hundred similarity matrices generated from data where band and measure labels were permuted within participants. We find similar correlations to the frequency band mask for both the spread and presence of IEDs (**Fig. 4C**; Pearson’s correlation (*n* = 3) *r*_*presence*_(112) = 0.35, *r*_*spread*_(112) = 0.30), although the correlation is slightly stronger for the presence of IEDs. Both correlations were stronger than all one-hundred correlations obtained from permuted null models (**Fig. S14**). To test our second hypothesis, we repeated the same analysis, but with a measure mask. Here, we find that correlations are strongest for the spread-of-IEDs predictor (**Fig. 4C**; Pearson’s correlation *r*_*presence*_(112) = 0.26, *r*_*spread*_(112) = 0.38). Here, the correlation for both predictors was larger than all one-hundred null models (**Fig. S14**). To test our third hypothesis, we repeated the same analysis with a grouped frequency (high or low) mask. Here, we find much larger correlations in the presence of IEDs (**Fig. 4C**; Pearson’s correlation *r*_*presence*_(112) = 0.59, *r*_*spread*_(112) = 0.37). Additionally, this mask explained much more variance than each band individually for the presence, but not spread, of IEDs. Here, the correlation for both predictors was larger than one-hundred null models (**Fig. S14**). Ultimately, we find that both the bands and measures tend to have more similar effects than would be expected by chance. Additionally, the presence of IEDs seems to show more similar changes to functional connectivity within bands than within measures, although those effects differ between high and low frequencies.

### Explanatory sources of individual variability

We next sought to understand sources of individual variability in changes to connectivity. However, participants in this study have coverage over a unique and heterogeneous set of regions, making it difficult to determine whether differences across individuals are due to differences in coverage or in characteristics of the participants. To test whether changes across participants may be due to differences in electrode placement, we wished to quantify whether connectivity changes were more similar within individuals or within regions. To quantify similarity, we calculated the Spearman’s correlation between the vector of connectivity changes across all bands, measures, and predictors between each individual contact (**Fig. 5AB**). We then calculated the correlation between these similarity matrices and masks selecting for entries from either the same individual or the same region. Here, regions are clinician-provided labels for each contact. We find a much larger Pearson’s *r* value for the participant mask, compared to the regions mask (Pearson’s correlation *r*_*part*_(2, 401, 318) = 0.13, *r*_*region*_(2, 401, 318) = 2.21 × 10^−3^). The low correlation value obtained from the participant level suggests that even features unique to each participant might have little explanatory power. However, the stark difference between participant and regional level analysis motivated us to continue investigating individual, rather than regional, differences in changes to connectivity resulting from IEDs.

We next tested which participant-specific features would best predict changes to connectivity for each band and measure using group LASSO regression. Group LASSO applies a penalty to groups of coefficients in a regression equation, allowing coefficients that do not greatly increase the explanatory power of the model to be regularized to zero. A parameter *λ* scales the regularization, such that a larger *λ* will result in a stricter penalty. Here, we chose the value of *λ* that minimized mean squared error across leave-one-out cross validation.

We hypothesized that clinical, contact, and demographic factors might impact the magnitude of IED effects on functional connectivity. We include four clinical variables: (1) presence or absence of lesion, (2) age at seizure onset, (3) the underlying etiology of epilepsy, and (4) the average rate of IEDs. We include five demographic variables: (1) age, (2) race, (3) sex, (4) handedness, and (5) institution. Lastly, we include three contact variables: (1) the cognitive system[38] that contains the contact with the most IEDs, (2) the hemisphere that contains the contact with the most IEDs, and (3) whether the participant had grid contacts, depth contacts, or both. Fifty-eight participants had all these fields of information.

We restrict our analyses to only measures that had changes in connectivity significantly different from zero in both IED detectors: the orthogonal amplitude envelope correlation, and imaginary coherence. The variables and corresponding regularized coefficients included in each grouped LASSO regression are shown in **Fig. 5C**. For categorical variables, the largest beta value assigned to a contrast at a given level was used. For both the presence and spread of IEDs, we found that some bands and measures were not well explained by any of the included variables. However, all variables except for sex and presence of lesion were included at least once. The most commonly included variable across both predictors was the rate of IEDs. This variable was included in five out of nine models that retained variables. Additionally, we found that the locus of IEDs (3/9), the underlying etiology (2/9), handedness (3/9), and institution are common across both IED features (2/9). We also noted some differences between the models that explain changes in the presence versus spread of IEDs. Age of seizure onset (2/9) was a commonly included variable for the presence of IEDs, and the hemisphere of IEDs (2/9) was common for the spread.

## DISCUSSION

Here, we conduct a thorough quantification of the changes in functional connectivity associated with IEDs in a large sample of 145 individuals with drug-refractory focal epilepsy. We test five different frequency bands and the broadband signal for changes associated with five different measures of functional connectivity and with three properties of IEDs. One key insight from our study is that, despite the tremendous heterogeneity in IED properties both within and between individuals[7, 61], we observe consistent increases in functional connectivity associated with IEDs. We find that these increases are not limited to regions in the irritative zone where IEDs occur, demonstrating that these increases are not a mere statistical property of the spikes themselves. In fact, we found that no subset of the tested edges robustly explained the increases observed in the global summary statistics. While it remains possible that specific edges may differentially influence the global increases we observe, it is notable that such increases are not limited to any of the edge subsets we tested.

Our study allows us to determine which properties of IEDs are responsible for the documented changes. We find that, once a single IED has occurred, more IED sequences within a time window do not further disrupt ongoing global functional connectivity, whereas IEDs that spread to more contacts further increase connectivity. Additionally, individuals whose functional connectivity changed significantly in response to the presence of an IED tended to show large changes associated with the spread of IEDs. Our work reveals that only amplitude-influenced measures (i.e. orthogonal amplitude envelope correlations and imaginary coherence) show changes, whereas phase-based and broadband measures tend to remain more stable, possibly due to the inconsistent presence of oscillations above the aperiodic background in our data. Across bands, we see that effects tend to be similarly sized within low and high frequency groups individually, but not across them. We also observe a high degree of individual variability in changes to functional connectivity associated with IEDs. No single feature robustly explained the variation observed across all measures and bands, but the rate of IEDs and the cognitive system that IEDs originate from explains the most variance. The changes reported here contribute substantial evidence towards an important unanswered question in epilepsy research, regarding the impact of IEDs on functional connectivity. Therefore, these results have important implications for preprocessing, analysis, and interpretation of research conducted with human iEEG data.

### IEDs disrupt multiple physiological processes

Our results show that responses to the presence of IEDs are more similar within groups of high and low frequencies than with each band individually. It is important to note that while this similarity structure is robust, the similarity between any pair of responses is often low, consistent with many studies showing frquency band specific changes in epilepsy or durng epileptic activity[22, 24]. While we cannot infer the mechanisms that generate oscillations from their frequency bands alone, previous research has reported differing roles for oscillations in low- versus high-frequency bands. For example, early intracranial recordings show that lower frequency oscillations (*<*30 Hz) tend to be much more spatially distributed than higher frequency oscillations[33, 62]. Further work has shown that power in the highest frequency band investigated here, high gamma, can be correlated with localized spiking activity[32, 34]. Additionally, while low frequency signals are thought to influence top-down cognitive processes, high frequency signals are thought to reflect bottom-up sensory driven processing[63, 64]. Here, we provide evidence that each of these processes is disrupted during IEDs, but might be impacted differently.

Our results point to two important future directions. First, it would be of interest to investigate changes in cross-frequency interactions and the slope of the power spectral density associated with IEDs. High and low frequencies have also been shown to have a preferential interaction direction, where the amplitude of higher frequencies is modulated by the phase of lower frequencies[30]. While these interactions can be spurious and require strict testing for the presence of oscillations above the 1/f background and against null models that preserve autocorrelation [65, 66], a detailed investigation of which of these interactions may change during IEDs would help further elucidate the neurophysiological processes that they disrupt. Second, opposing changes in high- and low-frequency bands can sometimes be explained by changes in the slope of the 1/f background, rather than changes in each band individually[67]. In studies of neural activation, the slope and power in specific bands are directly and intuitively related. However, it is unclear if such a relationship exists for connectivity. Additionally, here we do not see strongly anti-correlated patterns between low and high frequencies, and therefore we do not think this effect is driving the observed patterns. However, investigating changes in the slope of the 1/f background, as well as other non-sinusoidal features of the signal, would also add significantly to our understanding of how IEDs impact ongoing physiology.

We also show a separation between phase- and amplitude-based functional connectivity measures. Phase-based measures do not show significant connectivity changes that are consistent across detectors, and when changes are seen in individual detectors, they tend to be negative. In neuroscience, there is no definitive list of neural processes that can lead to changes in each space. However, changes to amplitude envelopes have been reported from bursting or sustained oscillations, as well as from synaptic potentials and spiking, whereas phase changes tend to be more often linked to oscillations. Therefore, the lack of consistent changes to functional connectivity might be attributed to the relatively small and variable proportion of contacts that tended to have oscillations in any given time window. While it is mathematically possible to have meaningful changes in phase synchrony between signals without prominent oscillations[68], we do not observe consistent changes in these processes during IEDs. Further investigation of oscillatory signals could help elucidate whether phase disruptions are a common feature of neural activity during IEDs.

### Implications for iEEG researchers

Our work has important implications for researchers who use iEEG data to understand basic features of the human brain that are independent of epilepsy and cannot be answered with other methods[69]. There exist multiple methods for discounting the impact of IEDs on this type of research, including removing some time points with IEDs, removing channels with IEDS, or including IED data. Here, we observe statistically significant increases in functional connectivity in a sample of individuals an order of magnitude larger than a typical iEEG study. The large-group effects extended beyond the contacts that showed IEDs. Additionally, some individuals showed large changes in functional connectivity associated with IEDs. Given that iEEG research is well-suited to within-individual experimental designs, we recommend, based on our results, that researchers consider removing entire time points that contain notable IEDs. We acknowledge that some individuals will have very high rates of IEDs, which could make this kind of methodological choice prohibitive, in which case the decision not to remove time points because of a high IED rate should be transparent in the methods. This approach will likely make results obtained from this population more consistent with the noninvasive EEG and fMRI literature in non-epileptic populations, and will likely lead to more robust inferences about generalized processes not linked to epilepsy.

Secondly, we also add to existing evidence that the choice of IED detector can matter for iEEG researchers[45, 48]. While our main findings are consistent across detectors, we do find the results across detectors are not identical. If IED detection is central to the claims of a paper, using multiple detectors and justifying the choices of detectors used could increase confidence in presented results.

### Unexplained variance in connections and individuals

Here, we test a few simple hypotheses about which connections might be contributing to increases in functional connectivity. Specifically, we test if increases in functional connectivity are preferentially explained by (1) the weakest edges in the distribution, (2) connections in the seizure onset zone, (3) between regions containing IEDs, (4) between grid and depth contacts, and (5) between grey and white matter contacts on depth electrodes.

We find that stark amount of the variance in responses is not explained by any measures tested here. The most consistent feature identified was that functional connectivity within pathological tissue in the SOZ, is increased more when IEDs spread to more contacts. The observed overlap between the two groups of pathological tissue areas is consistent with previous work, which showed that the functional group of spiking contacts overlaps with the SOZ is also the group with the highest IED rate, and that is responsible for most IEDs[6]. The largest effects are seen in higher frequency bands, which are thought to reflect more local activity. This pattern of findings is intuitive, given that the IEDs likely spread to regions within that pathological tissue. While edges outside these pathological regions still increase with broader spread of IEDs, this evidence suggests that the spread might be impacting more spatially local networks. However, none of the edge subsets we explored showed larger connectivity increases for the presence of IEDs. This observation is particularly noteworthy for grey and white matter contacts within depth electrodes. White matter has sometimes been considered to not have much meaningful signal, and be more appropriately used as a reference for recordings from grey matter[70]. However, here we see similar changes to functional connectivity in metrics that attempt to control for volume conducted signal, or any shared variance introduced through rereferencing. This is consistent with work suggesting that these contacts might be recording some signals from distant grey matter not recorded elsewhere[71].

Interictal connectivity changes relative to healthy controls show complex patterns of increases and decreases across systems[72, 73]. Additionally, these changes are temporally dynamic, but culminate in broad increases in connectivity during seizure initiation[35]. Given that IEDs are often considered to be a simple instantiation of epileptic activity that leads to seizures[8], the specific connections contributing to increases likely vary depending on the specific seizure semiology features of the individual.

Given the large variability in changes in functional connectivity across individuals, we also sought to characterize which features of individual participants explained some of the observed variance. We find that the identity of the participant explains more variance in responses than region the recording was taking from, but once again that most of the variance remains unaccounted for by either factor. The most consistently included variable was the cognitive system that most IEDs originated from. The system from which IEDs originate might be capturing information regarding the specific location of the seizure onset zone of a given individual and the associated epilepsy syndrome, i.e. frontal or temporal lobe epilepsy. Disentangling the contributions of these two features presents an exciting direction for future work. While we did not have access to each participants epilepsy syndrome directly, we expect these different phenotypes to show different patterns, due to the diverse spatial distribution of pathology and connectivity changes from controls[72–74]. Additionally, theoretical work on the spread of perturbations in brain networks suggests different directions of travel and patterns for perturbations originating in different regions[75, 76]. It is thus likely that both clinical and generalizable neural features underlie the explanatory power of the cognitive system where the IED originated.

While it is unsurprising that no single variable captured individual variability well, it is worth considering that some band-measure combinations were not well-explained by any combination of variables. However, the cohort used in this study was not homogeneous, and contains several additional sources of interindividual differences that are not captured by our variables. For example, we included no information about the shape or frequency of IEDs, treatment plans while in the epilepsy monitoring unit, or the type of epilepsy. A meta-analysis of smaller studies providing this information might better be able to address individual variability in observed effects.

## Limitations

This study presented a thorough characterization of how global functional connectivity changes during IEDs, and simplified the space of possible biophysical interactions that are affected during an IED. However, the findings from this study should be interpreted in light of limitations in our approach and methods. First, we sought to characterize changes to the macroscale behavior of the system, quantified as global connectivity. However, this approach prevented us from identifying even robust local changes in connections that counteract the shift in mean connection weight. Second, we chose to investigate a window size of 1 second, but it is possible that other time windows could capture complementary changes across different frequency bands. Third, we used a permutation-based linear model to estimate effect sizes, which precludes an identification of nonlinear effects associated with any of our predictors. Fourth, we tested for changes in the magnitude of functional connectivity metrics, but this approach creates inconsistency for metrics that use negative values (AR model, amplitude envelope correlation) by preventing the identification of changes that shift from large positive to large negative values.

This work used an extensive publicly available dataset in order to address questions about epilepsy and iEEG data in larger than standard cohorts. However, some of our analyses needed to carve out smaller groups from this large cohort. Most notably, smaller sample sizes were used for analyses of differences between grey and white matter and across etiologies. These analyses in particular would benefit from replications in independent datasets. One important final consideration is the potential for filtering artifacts to influence our results[42]. Filters, like those used when bandpass filtering data, can induce spurious oscillations and estimates of connectivity when applied to sharp transient activity or steps. For transparency, we show the results of applying the filters used in our preprocessing steps in **Fig. S2**. We are confident that our results are not strongly influenced by this type of artifact for the following reasons: (1) the lack of strong changes in phase-locking value associated with

IEDs, (2) larger changes in connectivity for contacts without IEDs, and (3) the significant differences seen in functional connectivity only from contacts with no IEDs. However, it is still possible that these artifacts could impact our findings in small ways.

### Conclusions and future directions

In addition to future studies investigating other aspects of neural activity, this exploratory work facilitates several avenues of hypothesis-driven research. While we were explicitly looking for large effects that could be visible after stringent multiple comparisons correction, specific hypotheses about more homogeneous groups that are subsets of this population could elucidate more subtle changes. For example, individuals with frontal versus temporal lobe epilepsy often have changes to functional connectivity in different systems at rest[72, 73]. Future work could test whether the presence or spread of IEDs impacts interactions between those systems selectively, or whether changes are only seen in resting-state activity. Additionally, while work has already shown a relationship between the presence of IEDs and task performance, an extension of this work to task data could assess whether that relationship extends to other features of IEDs such as their number and spread. Lastly, specific etiologies can have more homogeneous neural abnormalities. It would be interesting to investigate whether deviations from the average profile presented here are localized to regions of pathology.

A second exciting extension of this project involves testing how the changes to functional connectivity associated with IEDs might depend upon the current state of the brain. Other perturbation techniques, such as TMS, elicit state-dependent changes in neural activity and behavior [77]. Characterizing the state of the network preceding IEDs and assessing if these states modulate the response would provide a more complete understanding of the global response of the epileptic brain to IEDs.

In this work, we demonstrate consistent increases in amplitude-influenced functional connectivity associated with IEDs. We present evidence that IEDs influence multiple features of neurophysiology and that the observed changes in functional connectivity are not limited to pathological tissue. Based on these observations, we recommend that basic scientists working with iEEG data who wish to make claims about the general, non-epileptic population using functional connectivity, remove time points including IEDs from their analysis. Additionally, these observations demonstrate that the epileptic brain displays a tendency towards synchrony in the context of small perturbations, even when that synchrony does not amount to a seizure. Future work investigating and modeling the dynamics of seizures can now use these observations to better understand the principles underlying seizure generation.

## CITATION DIVERSITY STATEMENT

Recent work in several fields of science has identified a bias in citation practices such that papers from women and other minority scholars are under-cited relative to the number of such papers in the field [78–82]. Here we sought to proactively consider choosing references that reflect the diversity of the field in thought, form of contribution, gender, race, ethnicity, and other factors. First, we obtained the predicted gender of the first and last author of each reference by using databases that store the probability of a first name being carried by a woman [82, 83]. By this measure (and excluding self-citations to the first and last authors of our current paper), our references contain 7.04% woman(first)/woman(last), 14.08% man/woman, 18.31% woman/man, and 59.15% man/man. This method is limited in that a) names, pronouns, and social media profiles used to construct the databases may not, in every case, be indicative of gender identity and b) it cannot account for intersex, non-binary, or transgender people. Second, we obtained predicted racial/ethnic category of the first and last author of each reference by databases that store the probability of a first and last name being carried by an author of color [84**?**]. By this measure (and excluding self-citations), our references contain 9.14% author of color (first)/author of color(last), 12.9% white author/author of color, 24.4% author of color/white author, and 53.56% white author/white author. This method is limited in that a) names and Florida Voter Data to make the predictions may not be indicative of racial/ethnic identity, and b) it cannot account for Indigenous and mixed-race authors, or those who may face differential biases due to the ambiguous racialization or ethnicization of their names. We look forward to future work that could help us to better understand how to support equitable practices in science.

## ACKNOWLEDGEMENTS

We would like to thank Andrei Klishin for helpful discussions of the assumptions of cross-spectral analysis. D.S.B. and J.S. acknowledge support from the John D. and Catherine T. MacArthur Foundation, the Alfred P. Sloan Foundation, the ISI Foundation, the Paul Allen Foundation, the Army Research Laboratory (W911NF-10-2-0022), the Army Research Office (Bassett-W911NF-14-1-0679, Grafton-W911NF-16-1-0474, DCIST-W911NF-17-2-0181), the Office of Naval Research, the National Institute of Mental Health (2-R01-DC-009209-11, R01 – MH112847, R01-MH107235, R21-M MH-106799), the National Institute of Child Health and Human Development (1R01HD086888-01), National Institute of Neurological Disorders and Stroke (R01 NS099348), and the National Science Foundation (BCS-1631550, IIS-1926757). The content is solely the responsibility of the authors and does not necessarily represent the official views of any of the funding agencies.

