## Supplemental Materials for "Fluctuations in functional connectivity associated with interictal epileptiform discharges (IEDs) in intracranial EEG"

### Supplementary Materials for "Fluctuations in functional connectivity associated with interictal epileptiform discharges (IEDs) in intracranial EEG"

Jennifer Stiso[1,2], Lorenzo Caciagli[2], Peter Hadar[3], Kathryn A. Davis[3], Timothy H. Lucas[3], and Danielle S. Bassett[2,4,5,6,7,8,9]

[1] *Neuroscience Graduate Group, Perelman School of Medicine, University of Pennsylvania, Philadelphia, PA 19104, USA*
[2] *Department of Bioengineering, School of Engineering and Applied Science, University of Pennsylvania, Philadelphia, PA 19104, USA*
[3] *Department of Neurology, Perelman School of Medicine, University of Pennsylvania, Philadelphia, PA 19104, USA*
[4] *Department of Electrical & Systems Engineering, School of Engineering & Applied Science, University of Pennsylvania, Philadelphia, PA 19104, USA*
[5] *Department of Psychiatry, Perelman School of Medicine, University of Pennsylvania, Philadelphia, PA 19104, USA*
[6] *Department of Physics & Astronomy, College of Arts & Sciences, University of Pennsylvania, Philadelphia, PA 19104, USA*
[7] *The Santa Fe Institute, Santa Fe, NM 87501, USA and*

#### SUPPLEMENTAL ANALYSES

##### Contribution of the aperiodic component of signals

Some recent studies that investigate power in specific frequency bands separate the aperiodic backbone of the signal's power spectral density (PSD) from true oscillations which arise above that backbone. This separation has been fruitful, and allowed researchers to decipher different potential mechanisms for observed phenomena. However, precisely how the oscillatory and aperiodic components of a signal impact connectivity estimates from instantaneous amplitude and phase is less well understood, and warrants further investigation in future work. Although a full investigation is not within the scope of our paper, we nevertheless wished to demonstrate the extent to which our results could be driven selectively by the aperiodic component of the signal, especially given our decision to control for spectral power in our regression analyses.

We formulated 4 hypotheses about how the aperiodic component might affect our results: (1) regions with more similar slopes of the aperiodic component would have stronger functional connectivity; (2) regions with more similar slopes and oscillations will have higher functional connectivity; (3) pairs of regions with steeper slopes would have stronger functional connectivity; (4) pairs of regions with steeper slopes and oscillations will have stronger functional connectivity. To test these 4 hypotheses, we selected two channels from our datasets, and altered the power spectral densities to match each of these four conditions. We then used linear regression to calculate the effect of the slope of the aperiodic component on functional connectivity while controlling for spectral power.

Specifically, we used IRASA (irregular re-sampling auto-spectral analysis) to estimate the power spectral density and aperiodic component of two channels (channel A and channel B). To test hypothesis 1, we then aligned the aperiodic component of channel B to that of channel A, and generated 50 additional PSDs where the slope of the aperiodic component of channel B got incrementally further from channel A. We then generated 50 timeseries with random phase instantiations for each PSD, and calculated the effect of slope on functional connectivity for all phases. To test hypothesis 2, we repeated the same analysis, but added an oscillation at 20 Hz to the PSD of channel B. To test hypothesis 3, the PSDs of channels A and B were aligned, and then both channels' slopes were increased incrementally. The same method was used to test hypothesis 4, but oscillations at 20 Hz were also added to the PSDs.

If there was a strong relationship between the apreiodic component of the signal and functional connectivity, we would expect small $p$-values and large coefficients for some measures, and some tests. However, for all hypotheses, we find that $p$-values are mostly uniformly distributed between 0 and 1, and coefficients are uniformly distributed around 0 (**Fig. S6-9**).

##### Sources of individual variability

We first tested whether differences in electrode type mediated these individual differences. Some individuals in the cohort were implanted with only flat grid or strip electrodes on the surface of the brain, while others were implanted with only penetrating or depth electrodes that reached deeper structures. These two types of electrodes sample the brain differently, and can record from different neural structures. We hypothesized that these differences in sampling might lead to different magnitudes of changes to functional connectivity. However, we found that no band-measure combination showed a statistically significant main effect for electrode type after multiple comparisons correction (**Fig. S11**; permutation-based linear model).

Next, we tested the hypothesis that each individual's etiology would explain individual differences in connectivity. For all etiologies with at least 10 individuals (unknown, other, MCD, TBI, infection, other), we tested for a significant main effect of etiology on changes in functional connectivity with a permutation based linear model. We found no significant main effects after multiple comparisons corrections (**Fig. S18**).

The above analyses elucidate the extent to which individual differences in connectivity changes are explained by etiology. One can also ask whether our broad conclusions about which connectivity measures change during IEDs are similar in smaller, more homogeneous groups. To answer this question, we calculated the magnitude of changes in connectivity (as shown in **Fig. 1**) for only the TBI and MCD populations. These populations were selected because they are the largest, most homogeneous etiology categories. We test whether effect sizes in each group are similar to the full sample, rather than comparing $p$-values because of the smaller sample sizes in each group. We find that for the presence of IEDs, the TBI group has effect sizes more similar to the full population than the MCD group. For the spread of IEDs, both groups have highly similar effect sizes to our full population (Spearman's correlation: $\rho_{TBI,presence} = 0.67$, $p_{TBI,presence} = 6.8 \times 10^{-5}$, $\rho_{TBI,spread} = 0.66$, $p_{TBI,spread} = 1.2 \times 10^{-4}$, $\rho_{MCD,presence} = 0.46$, $p_{MCD,presence} = 0.01$, $\rho_{MCD,spread} = 0.81$, $p_{MCD,spread} = 1.07 \times 10^{-6}$). This analysis shows that most of our conclusions are highly similar across these two etiologies. However, the lower correlations for the presence of IEDs in individuals with MCD also suggests that there might be interesting differences in which measures change across groups.

Lastly, we tested the hypothesis that the average rate of IEDs would explain individual differences in connectivity (**Fig. S19**).

#### SUPPLEMENTAL FIGURES

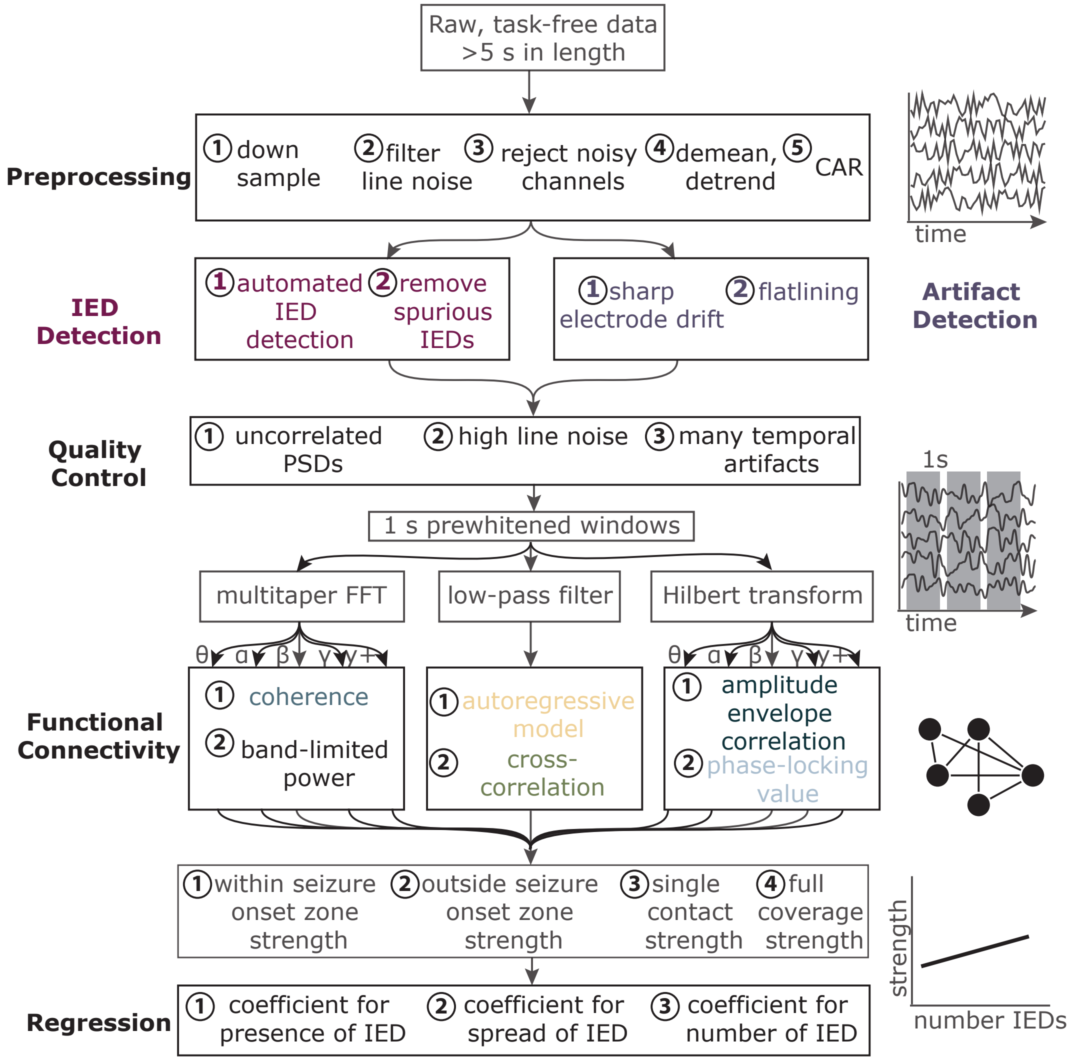


FIG. 1. **Flow Chart of Methods.** (Preprocessing) Epochs of at least 5 seconds undergo preprocessing steps 1-5. Data are downsampled to 500 Hz, and re-referenced to a common average (CAR). (IED Detection) Preprocessed data then undergo detection and selection of IEDs. (Artifact Detection) In the same preprocessed data, we detect transient artifacts, including sharp electrode drift and periods of flatlining. These epochs are removed from further analyses. (Quality Control) Entire datasets are rejected based on any of 3 criteria numbered in the box. (Functional Connectivity) Data are then segmented into 1-second windows, and prewhitened within each window. All windows then undergo three manipulations: a multitaper fast Fourier transform (FFT), low-pass filtering, and Hilbert transform. FFTs and Hilbert transforms are performed in 5 frequency bands. The multitaper FFT is then used to calculate coherence and power in each band. Low-pass filtered data is used to calculate a cross-correlation and autoregressive fit. Hilbert-transformed data is used to calculate an amplitude envelope correlation, and the phase-locking value. Functional connectivity measures are then summarized as the strength (mean connectivity) in 4 groups of channels: those in the seizure onset zone (SOZ), those outside the SOZ, each contact individually, and all channels. (Regression) Lastly, we fit a permutation-based linear model for each subject to obtain the standardized coefficients for a linear model associating a categorical indicator of an IED, the spread of IEDs, and the number of IEDs per window with each measure of strength. Power is included as a nuisance covariate in order to assess changes in functional connectivity that cannot be explained by changes in activity.

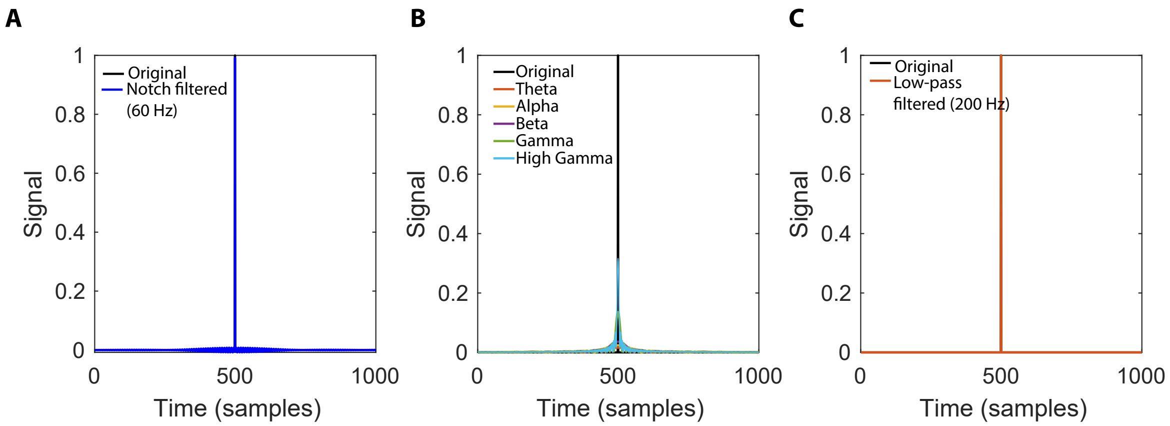


FIG. 2. **Filter responses.** *(A)* The result of notch filtering a single spike with the same parameters used in the main text. *(B,C)* The same as panel *(A)*, but for the bandpass and low-pass filters used in the main text.

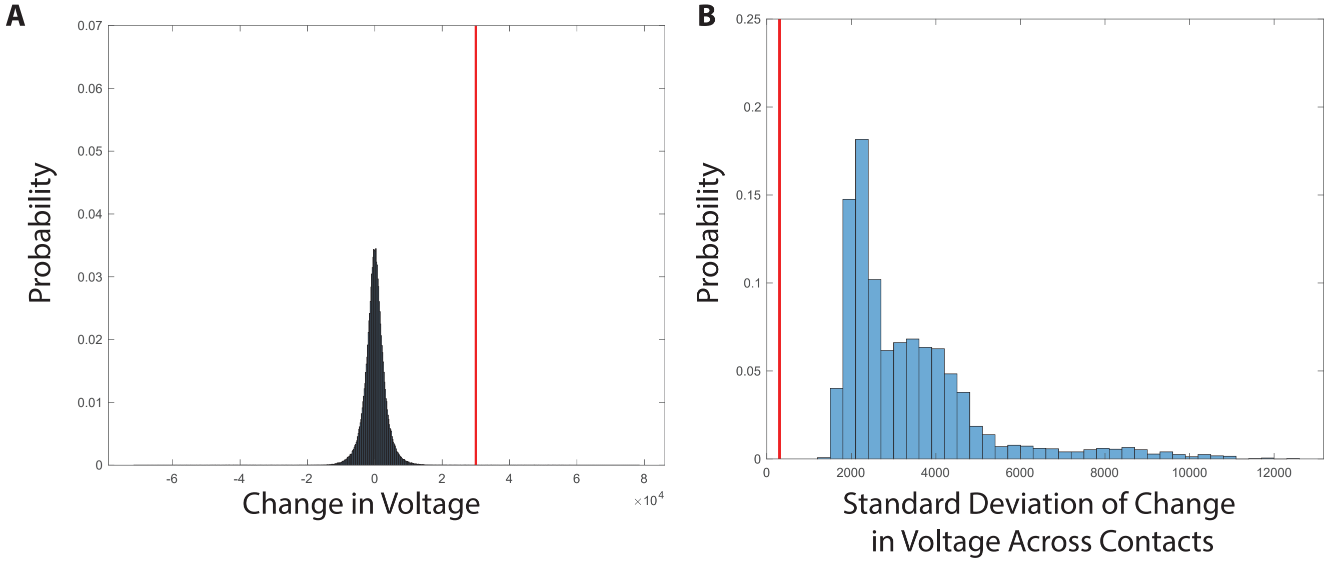


FIG. 3. **Thresholds for artifact detection.** *(A)* The change in voltage from 1 second of data randomly selected from each dataset. Red line indicates the threshold used to identify artifacts. The change in voltage was used to identify sharp transient artifacts present in timeseries. *(B)* The same as panel *(A)*, but for the variance in the change in voltage across contacts. This value was used to detect periods of flatlining in recordings.

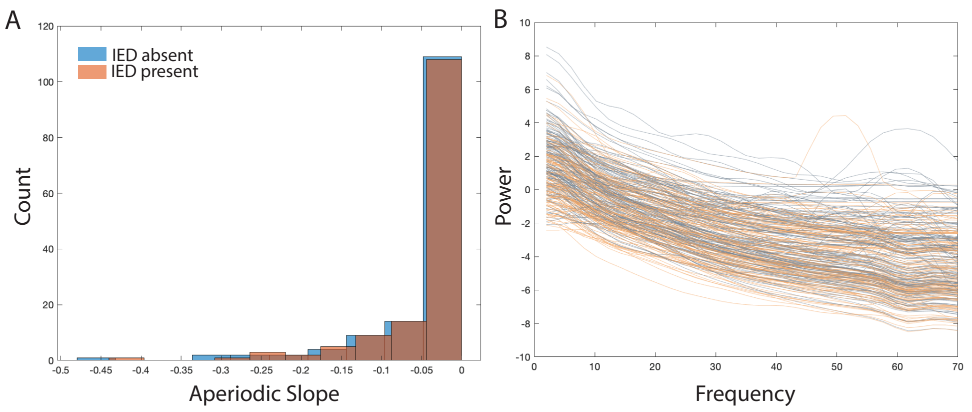


FIG. 4. **Power spectral densities for the presence and absence of IEDs.** ]*(A)* The slope of the aperiodic component of the power spectral density for each dataset. Slopes were averaged across windows and contacts. The blue histogram shows the slopes of windows without IEDs, whereas the orange histogram shows the slopes of windows with IEDs. *(B)* Power spectral densities for each dataset. Power spectral densities are averaged across windows and channels. Blue power spectral densities are from windows where IEDs are absent, whereas orange power spectral densities are from windows where IEDs are present.

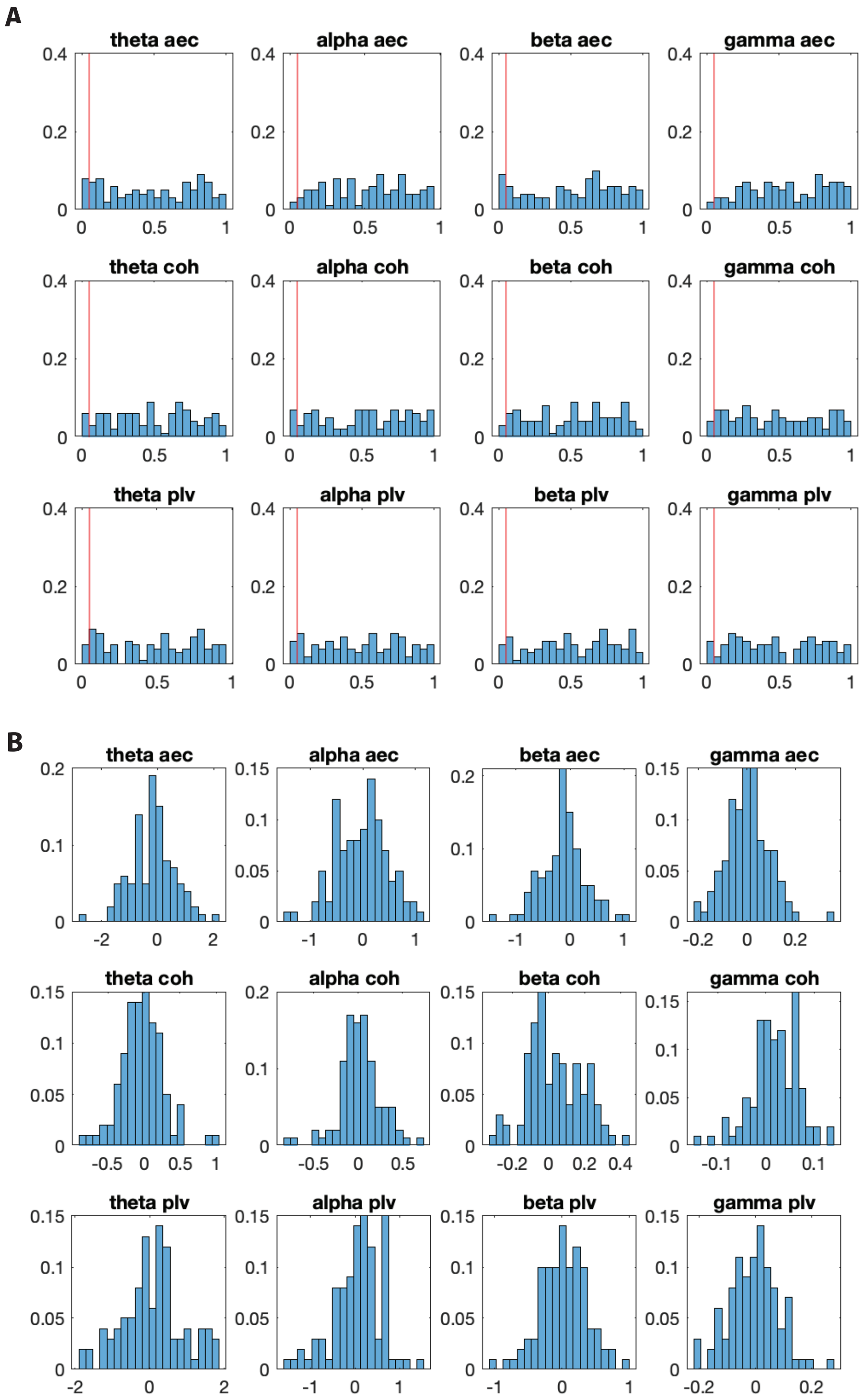


FIG. 5. **Influence of the aperiodic component of the signal on functional connectivity.** *(A)* The results of regression analyses between the similarity of the slope of the aperiodic component of two signals and the functional connectivity between those two signals, after controlling for spectral power. Distributions show values for different phase distributions of the signals. The $p$-values are shown on the left, with the red line indicating $p = 0.05$. *(B)* Shows the coefficients.

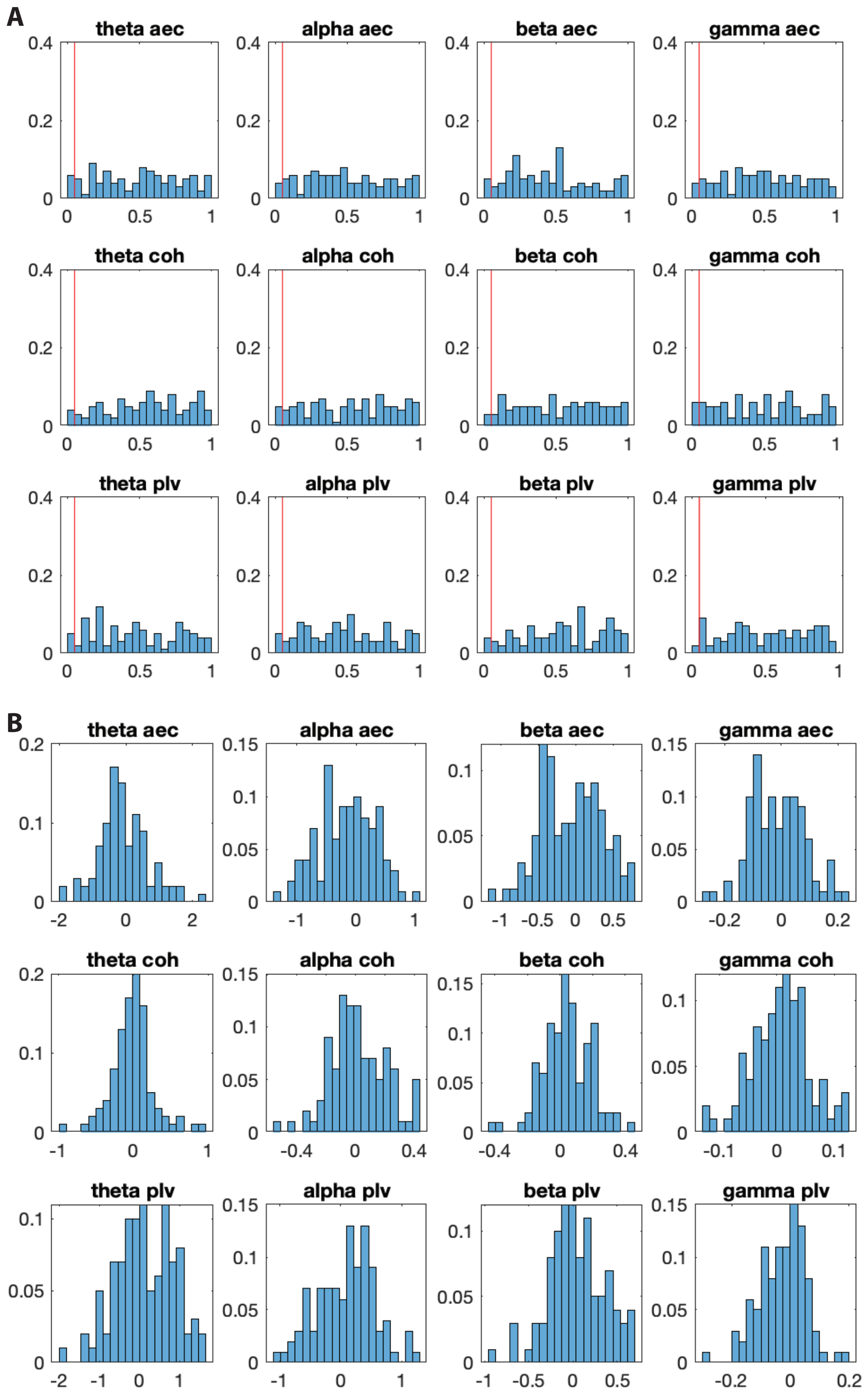


FIG. 6. **Influence of the aperiodic component of the signal on functional connectivity.** The results of regression analyses between the similarity of the slope of the aperiodic component of two signals, one with a 20 Hz oscillation present and the functional connectivity between those two signals, after controlling for spectral power. Distributions show values for different phase distributions of the signals. *(A)* Shows the $p$-values, with the red line indicating $p = 0.05$. *(B)* Shows the coefficients.

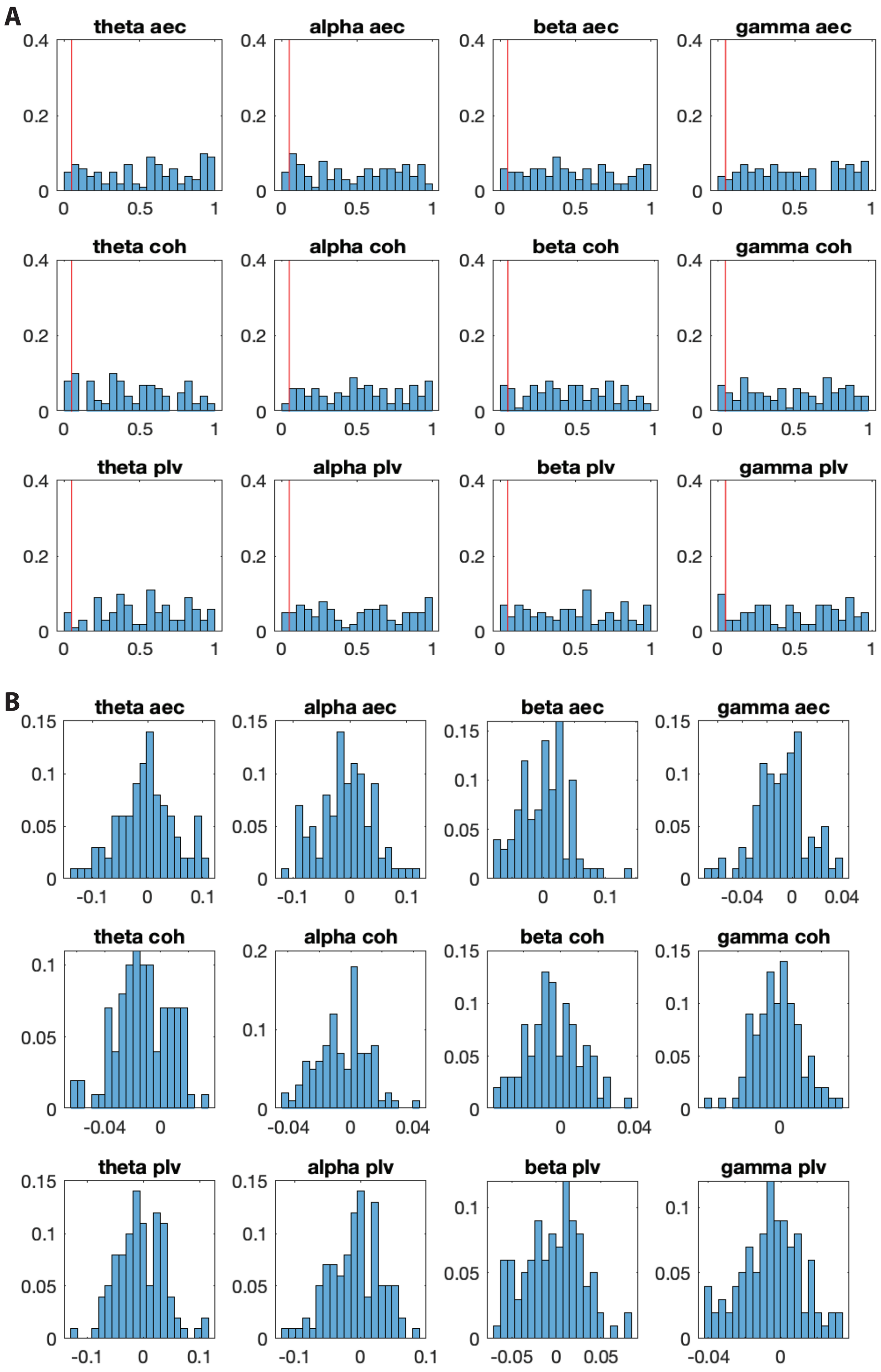


FIG. 7. **Influence of the aperiodic component of the signal on functional connectivity.** The results of a regression analyses between the steepness of the slope of the aperiodic component and the functional connectivity, after controlling for spectral power. Here, both channels always had the same slope, but the value of that slope differed. Distributions show values for different phase distributions of the signals. *(A)* Shows the $p$-values, with the red line indicating $p = 0.05$. *(B)* Shows the coefficients.

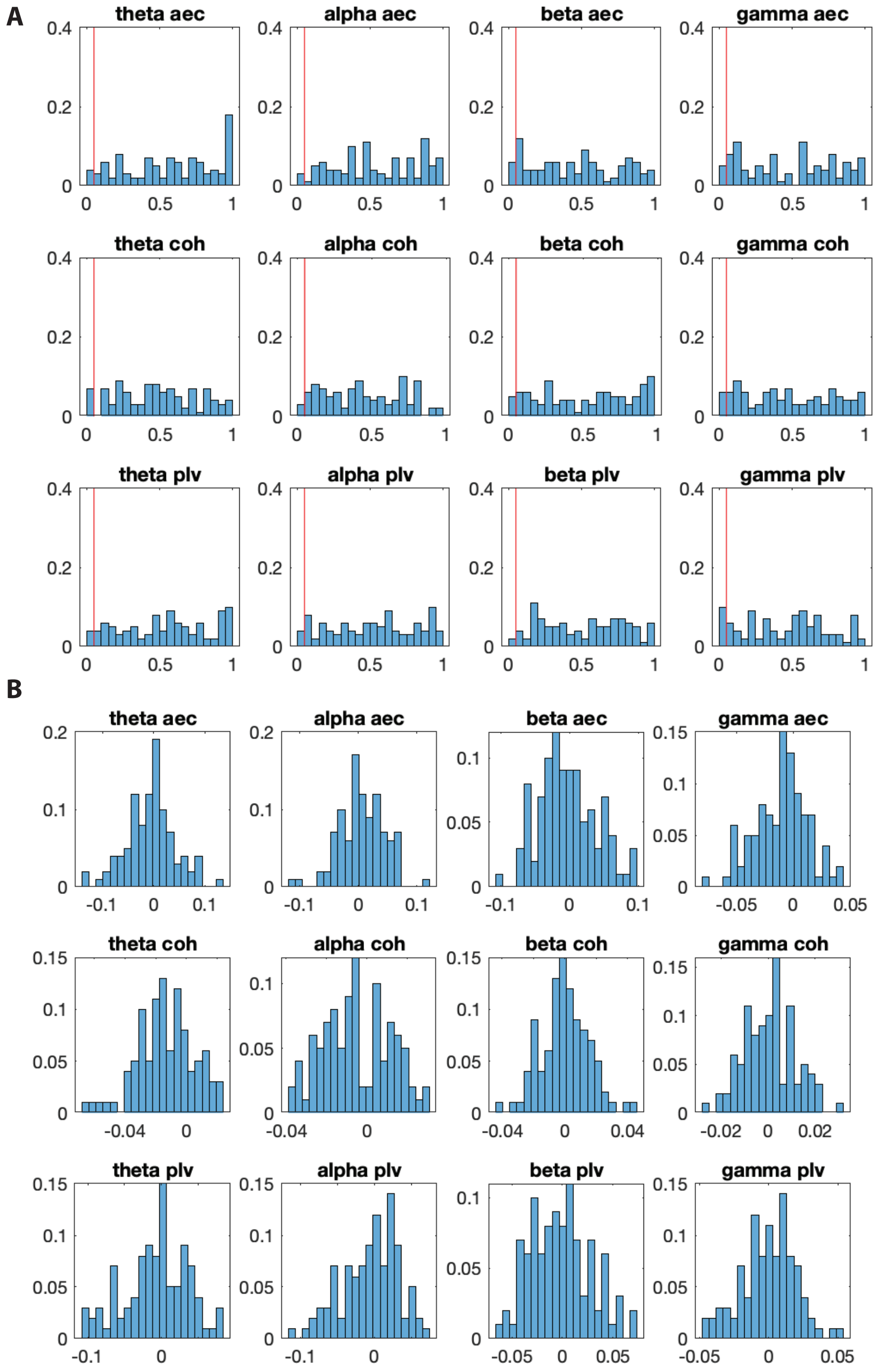


FIG. 8. **Influence of the aperiodic component of the signal on functional connectivity.** The results of regression analyses between the similarity of the slope of the aperiodic component of two signals with 20 Hz oscillations present and the functional connectivity between those two signals, after controlling for spectral power. Distributions show values for different phase distributions of the signals. *(A)* Shows the $p$-values, with the red line indicating $p = 0.05$. *(B)* Shows the coefficients.

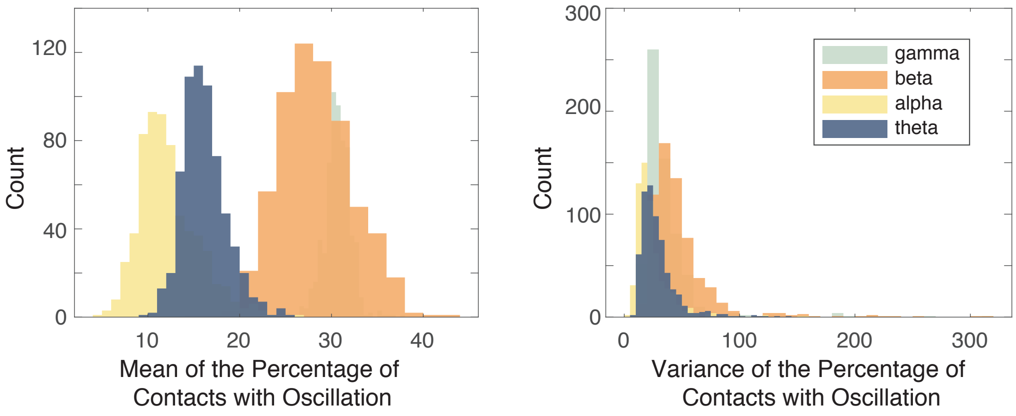


FIG. 9. **The prevalence of oscillations.** *(left)* The mean percentage of contacts with oscillations above the aperiodic background in each 1 second window. Since a network with $n$ nodes will have $n^2$ edges, we can use this information to infer the number of edges that would be between two nodes with oscillations. For example, a network with 30% of nodes containing oscillations would have 9% of its edges between nodes with oscillations. A network with 10% of its nodes containing oscillations would have 1% of its edges between nodes with oscillations. Different bands are shown in different colors. *(right)* The variance in the percentage of contacts with oscillations above the aperiodic background.

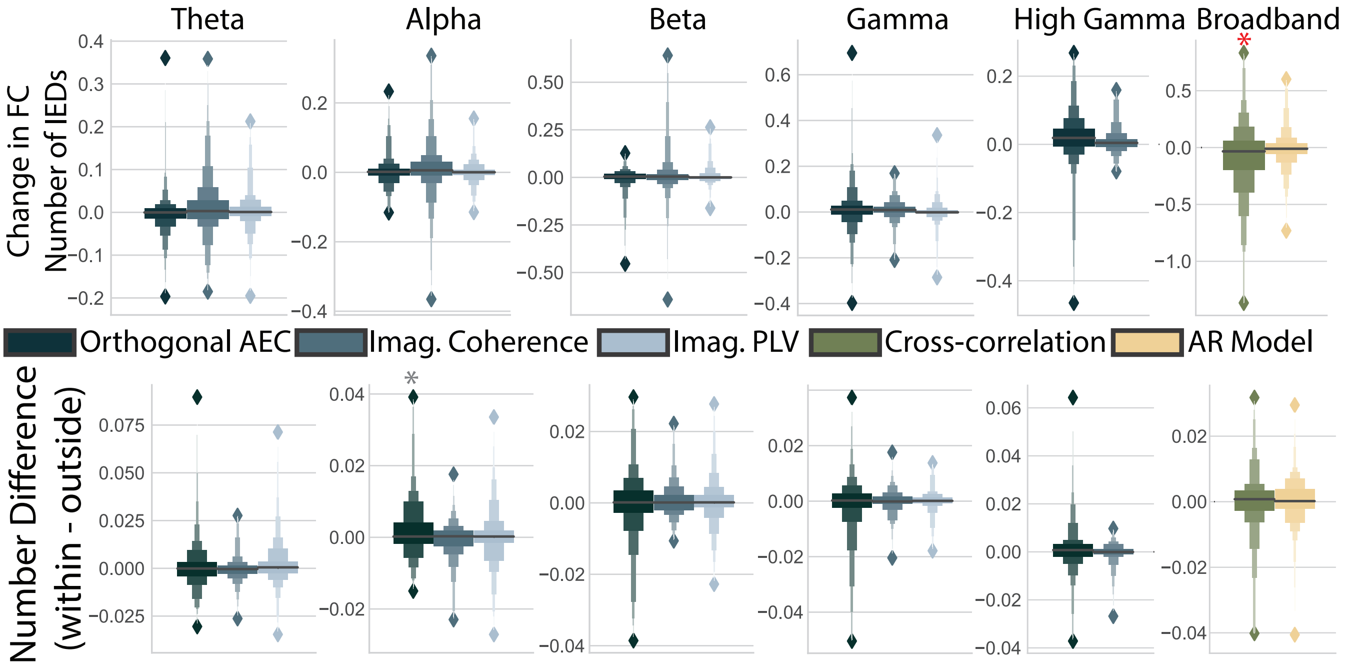


FIG. 10. **Differences in groups of connections for the number of IEDs.** *(Top)* Distributions of coefficients for changes in the skew of connections associated with the number of IEDs. Obtained from permutation-based regression including all IED predictors, power in a given band, and the recording session. Columns indicate different frequency bands, and colors indicate different measures. Bright red asterisks indicate significant distributions after multiple comparisons correction that were reproduced with a different spike detector. * = $p < 0.05$, ** = $p < 0.01$, *** = $p < 0.001$, **** = $p < 0.0001$. *(Bottom)* The same as in the top panel, but for the coefficients for connections outside the seizure onset zone subtracted from those within the seizure onset zone.

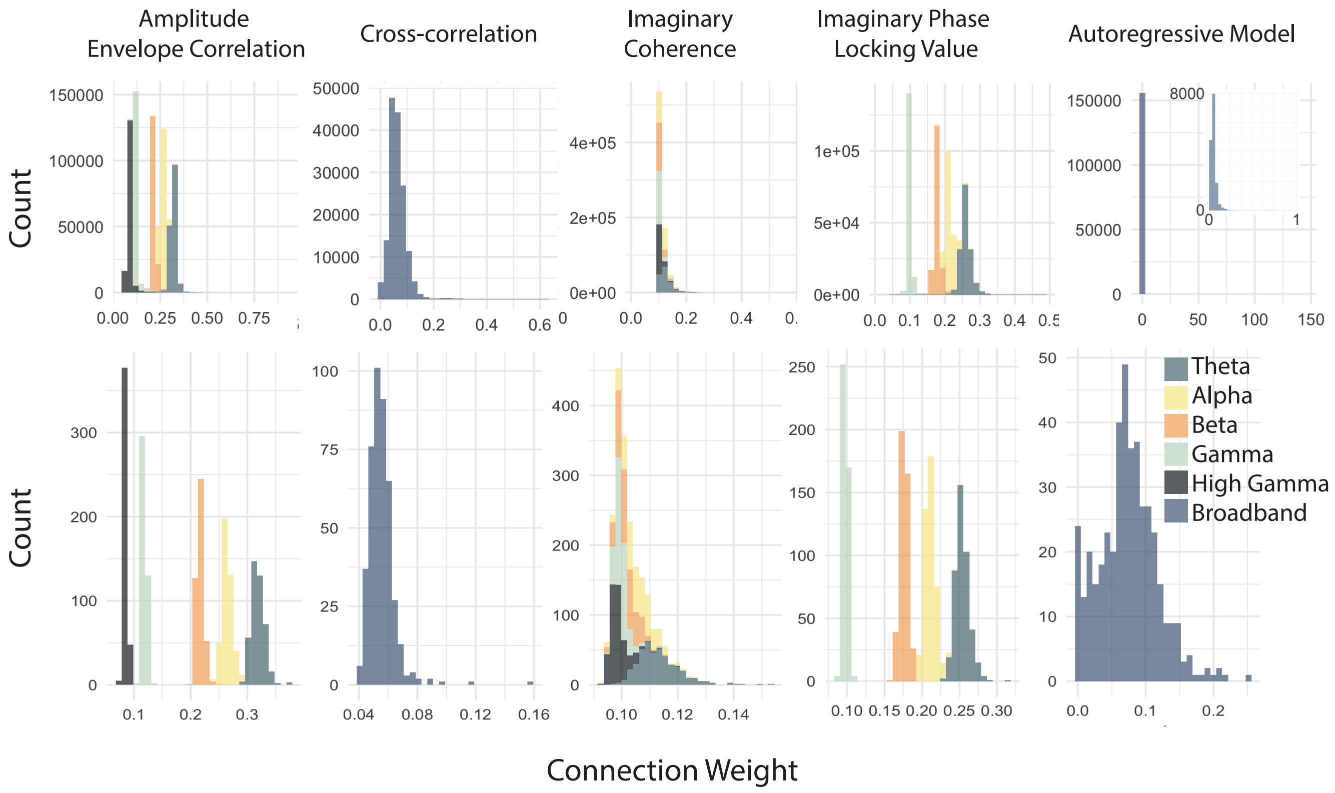


FIG. 11. **Strength distributions.** The distribution of strength values across connections and time points for all participants *(top)* and for the first participant in the dataset *(bottom)*.

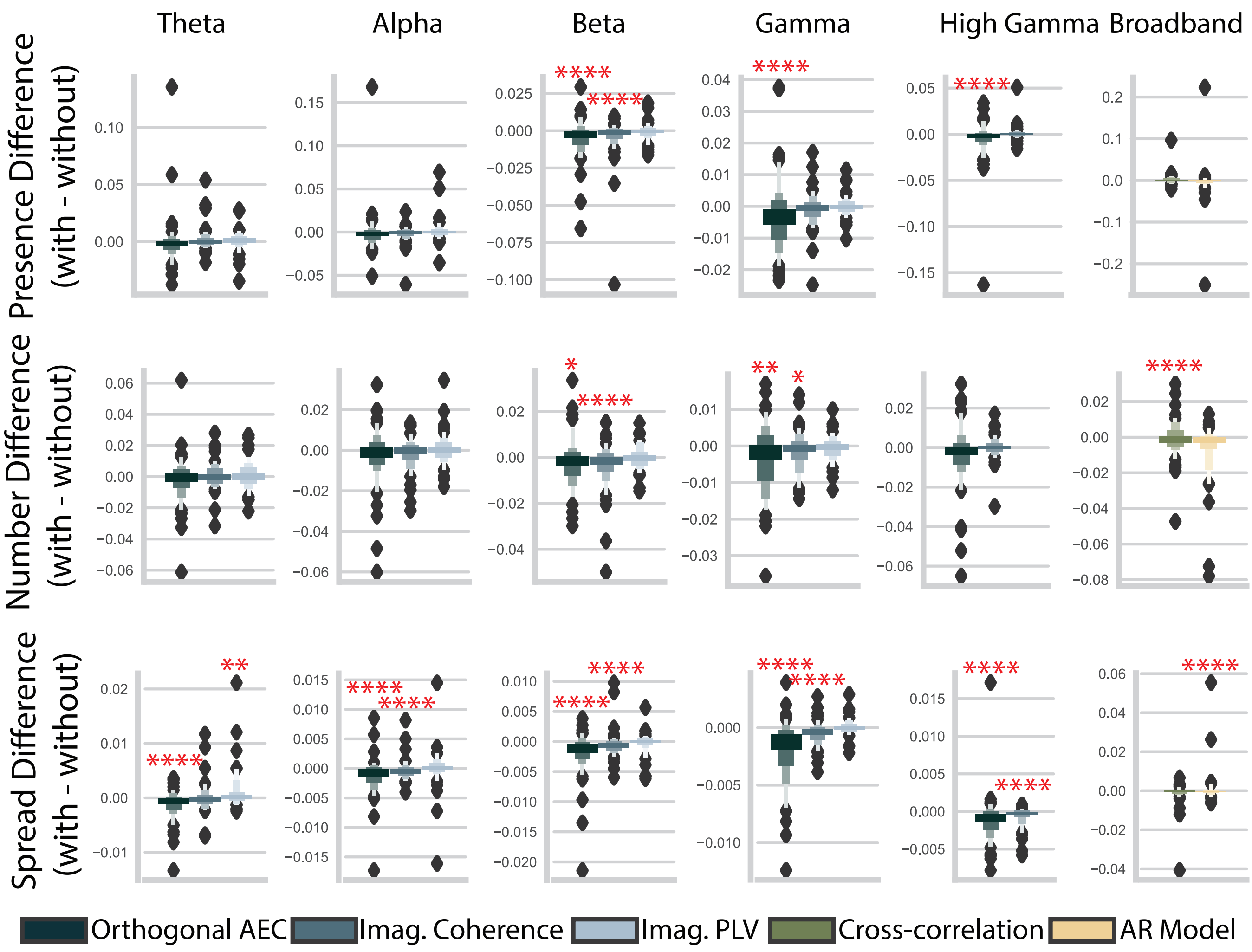


FIG. 12. **Differences in effect sizes between contacts with and without IEDs.** Distributions of coefficients for the presence *(top)*, number *(middle)*, and spread *(bottom)* of an IED. Columns indicate different frequency bands and colors indicate different measures. * = $p < 0.05$, ** = $p < 0.01$, *** = $p < 0.001$, **** = $p < 0.0001$.

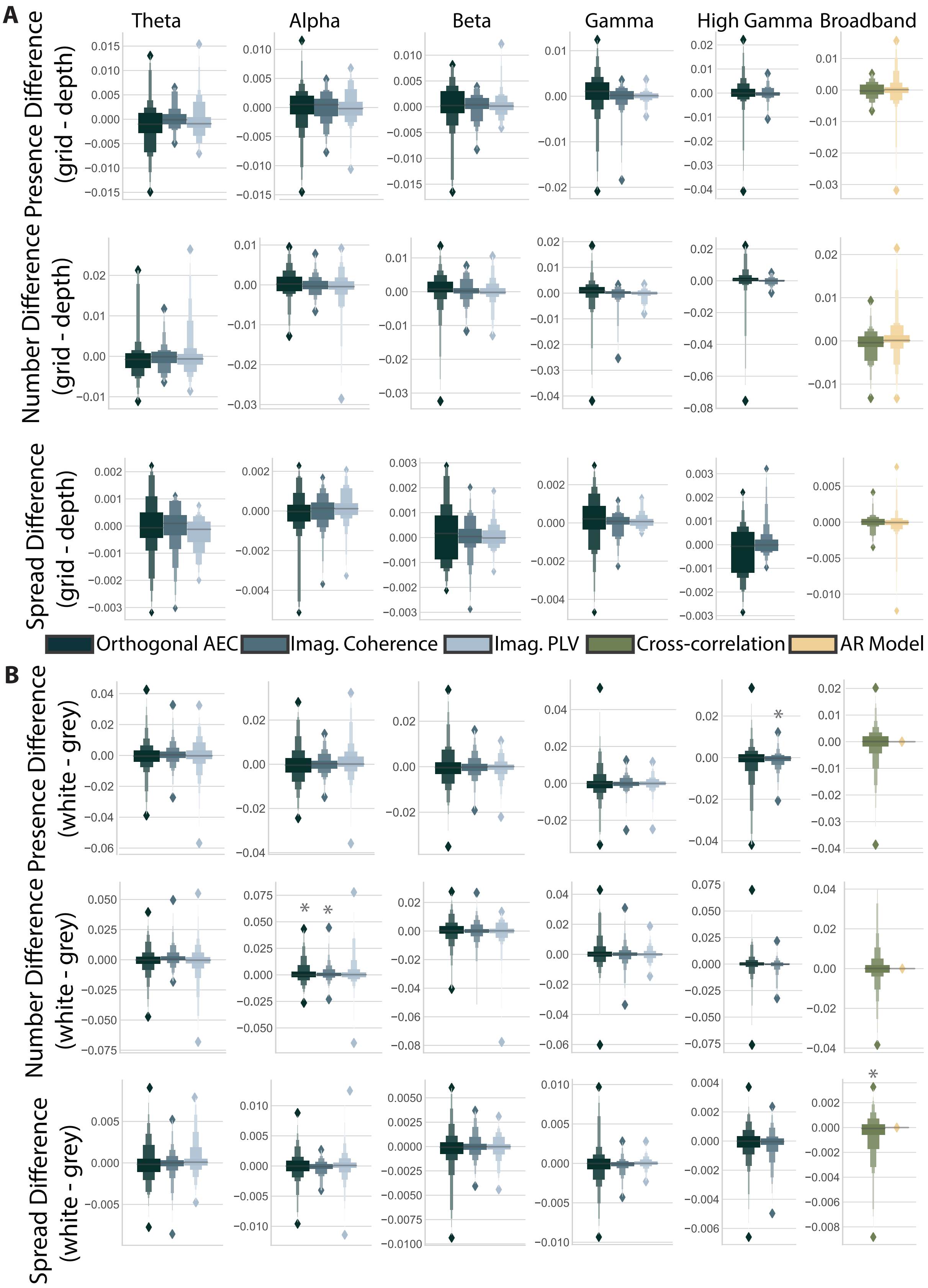


FIG. 13. **Differences in effect sizes between contacts within grid versus depth contacts, and within grey versus white matter.** *(A)* Distributions of the difference in functional connectivity for grid and depth contacts for the presence *(top)*, number *(middle)*, and spread *(bottom)* of an IED. Columns indicate different frequency bands, and colors indicate different measures. * = $p < 0.05$, ** = $p < 0.01$, *** = $p < 0.001$, **** = $p < 0.0001$. *(B)* The same as in panel *(A)*, but for the difference in the change in functional connectivity for white and grey matter contacts.

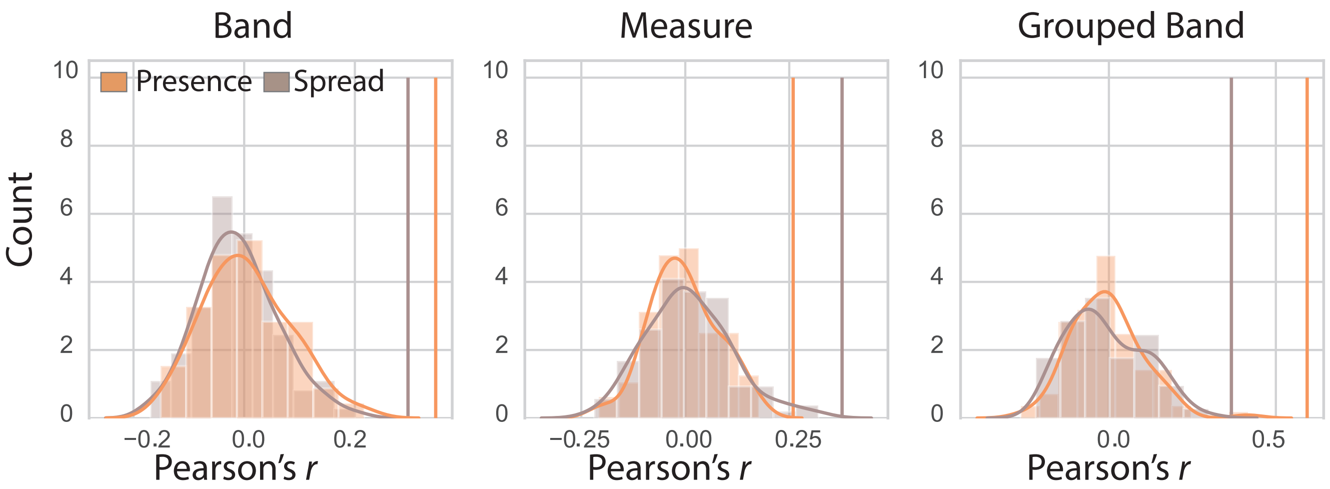


FIG. 14. **Permutation tests for similarity.** Correlations with different masks (band, measure, and grouped band) and 100 similarity matrices from permuted data from the presence of IED (orange) and spread of IED (brown) predictors. Kernel density estimate is shown in bold. Empirical correlation values are shown as vertical lines.
